# Polycomb repression during S/G2 phases restrain initiation of cell differentiation to the G1 phase of the cell cycle

**DOI:** 10.1101/2022.07.26.501502

**Authors:** Helena G. Asenjo, María Alcazar-Fabra, Mencía Espinosa, Lourdes Lopez-Onieva, Amador Gallardo, Emilia Dimitrova, Angelika Feldmann, Tomas Pachano, Jordi Martorell-Marugán, Pedro Carmona-Sáez, Antonio Sanchez-Pozo, Álvaro Rada-Iglesias, Robert J. Klose, David Landeira

**Author notes:** **Correspondence** to David Landeira.

## Abstract

The potential of pluripotent cells to respond to developmental cues and trigger cell differentiation is enhanced during the G1 phase of the cell cycle, but the molecular mechanisms involved are poorly understood. Variations in polycomb activity during interphase progression have been hypothesized to regulate the cell-cycle-phase-dependent transcriptional activation of differentiation genes during lineage transition in pluripotent cells. Here, we asked whether the Polycomb Repressive Complex 1 (PRC1) modulates the ability of mouse embryonic stem cells (ESCs) to differentially respond to developmental cues depending on the phase of the cell cycle in which they are found. We discovered that recruitment of PRC1 complexes and their associated molecular functions, ubiquitination of H2AK119 and three-dimensional chromatin interactions, are enhanced during S and G2 phases compared to the G1 phase. In agreement with the accumulation of PRC1 at target promoters upon G1 phase exit, cells in S and G2 phases show firmer transcriptional repression of developmental regulator genes that is drastically perturbed upon genetic ablation of the PRC1 catalytic subunit *Ring1b*. Importantly, depletion of Ring1b during retinoic acid stimulation interferes with the preference of mESCs to induce the transcriptional activation of differentiation genes in G1 phase. We propose that incremental enrolment of polycomb repressive activity during interphase progression reduces the tendency of cells to respond to developmental cues during S and G2 phases, facilitating activation of cell differentiation in the G1 phase of the pluripotent cell cycle.

## Introduction

Understanding the molecular basis of pluripotency is a fundamental challenge in developmental biology, and it is required for successful application of pluripotent cells to disease modelling and cell therapy ^1-3^. The response of individual cells within a given pluripotent cell population to differentiation signals can be diverse, in part as a consequence of the phase of the cell cycle in which they reside ^4,5^. The cell-cycle-phase-dependent response to cell differentiation is an evolutionary conserved mechanism in eukaryotes that is present in different stem cells from several organisms and tissues ^6-10^.

Despite its wide implications in developmental biology, the molecular pathways enabling stem cells to enter cell differentiation at a particular phase of the cell cycle remain to be characterized. In the case of pluripotent cells, initial observations suggesting that differentiation potential was interlocked with the regulation of cell cycle ^11^ were later confirmed by studies demonstrating that pluripotent cells in G1 phase are more prone to induce expression of developmental genes and effectively differentiate than cells in S and G2 phases ^12-16^. The higher tendency of G1 cells to exit pluripotency depends on the combined action of several mechanisms ^17^, of which current studies suggest that the transcription factor Smad2/3 and the chromatin proteins polycomb and trithorax might be key regulators ^13, 16, 18, 19^.

Polycomb Group (PcG) proteins are hallmark epigenetic regulators of development and cell differentiation in vertebrates ^20, 21^. PcG proteins associate to form different multimeric complexes known as Polycomb Repressive Complex 1 and 2 (PRC1 and PRC2) ^20, 21^. PRC1 complexes are defined by the presence of the catalytic subunit Ring1a/b, which mediate ubiquitination of lysine 119 on histone H2A (H2AK119ub1) ^22, 23^. Likewise, PRC2 complexes always contain Ezh1/2 proteins, which harbour the histone methyltransferase activity against lysine 27 on histone H3 (H3K27)^20, 24^. Importantly, PRCs form functionally specialized subcomplexes known as canonical PRC1 (cPRC1), variant PRC1 (vPRC1), PRC2.1 and PRC2.2 that include different accessory subunits ^20, 21^. Gene regulation by PRCs is achieved by the coordinated action of these functionally specialized subcomplexes that can lead to the formation of polycomb repressive chromatin domains that are highly enriched for H2AK119ub1 and H3K27me3 ^20, 21^. In mammalian pluripotent cells, PRCs repress the transcription of hundreds of lineage-specifier genes, implicated in pluripotent cell differentiation and early embryo development ^25^. Remarkably, virtually nothing is known about how the composite function of PRC subcomplexes is coordinated with chromatin changes occurring during cell cycle transition. This is probably an essential aspect of polycomb function because it intertwines with the essential property of pluripotent cells to maintain a delicate balance between preserving transcriptional memory during self-renewal and allowing the erasure of such memory during induction of cell differentiation. Notably, the PRC2 catalytic subunit Ezh2 is regulated by cyclin-dependent kinases 1 and 2 (CDK1 and CDK2) ^26-29^, and PRC2 subcomplexes are differentially recruited to target genes depending on the phase of the cell cycle ^19^, suggesting that mechanistic coupling of polycomb activity to the cell cycle machinery might be an unrecognized fundamental feature of the polycomb epigenetic system in eukaryotes.

In this study we have analysed whether PRC1 complexes regulate the ability of mouse embryonic stem cells (mESCs) to preferentially induce cell differentiation in the G1 phase of the cell cycle. We found that differential recruitment of PRC1 complexes at different phases of the cell cycle leads to the accumulation of cPRC1, vPRC1 and H2AK119ub1 during S and G2 phases. This is coupled to stronger promoter-promoter three-dimensional (3D) interactions and enhanced transcriptional repression of PRC1-bound genes. Importantly, depletion of Ring1b protein disturbs gene repression during S and G2 phases and hinders the cell-cycle-phase-dependent activation of PRC1 target genes upon induction of cell differentiation with retinoic acid. Overall, these data show that reduced activity of PRC1 complexes during G1 phase sets a chromatin state that facilitates the activation of developmental genes in response to differentiation cues in pluripotent cells.

## Results

### Ring1b remains bound to target gene promoters across interphase

We used previously published mESCs that express the fluorescent ubiquitination-based cell cycle indicator (FUCCI) reporter system (FUCCI-mESCs) to obtain highly enriched populations of cells in G1, S and G2 phases using flow cytometry (Figure S1A, B and C) ^19^ (Table S1). Western blot analysis of whole cell lysates and chromatin fractions showed that the PRC1 catalytic subunit Ring1b, the cPRC1-specific subunit Cbx7 and the vPRC1-specific subunit Rybp are expressed and bound to chromatin at similar amounts in G1, S and G2 phases (Figure S1D, E). Likewise, the level of H2AK119ub1 was also constant across interphase (Figure S1D, E). To address whether the distribution of PRC1 complexes on the genome changes during interphase progression, we compared the genome-wide binding profiles of the PRC1 catalytic subunit Ring1b in cells sorted in G1, S and G2 phases, using chromatin immunoprecipitation followed by sequencing (ChIP-seq). Ring1b was bound to 18.129 sites in the genome in S phase, with a clear tendency to be present at the promoter of genes at all phases of the cell cycle (Figure S1F, G). Ring1b was bound to similar number of gene promoters in G1 (n=7975), S (n=9322) and G2 (n=7422) phases, and these were largely overlapping (Figure S1H), indicating that there was no drastic reorganization of Ring1b binding during interphase transition. Notably, recruitment of Ring1b to target gene promoters (n=10908) (Figure S1H) was very similar in cells in G1 and G2 phases (Figure 1A, B, C and S1I). We detected a mild increased recruitment of Ring1b during S phase (Figure 1B), which might be related to the alternative role of PRC1 in DNA repair ^30-32^. As expected, Ring1b was bound to previously identified high confidence bivalent genes (HC bivalent) ^19^, which are targeted by PRC2 and are positive for both H3K4me3 and H3K27me3 (93.8%, 1575 out of 1678 genes) (Figure 1D). In fitting with the general trend detected at Ring1b target promoters (Figure 1B), HC bivalent genes showed similar binding profiles of Ring1b around the TSS in G1 and G2 phases (Figure 1E, S1J). To substantiate these observations, we repeated the Ring1b ChIP-seq using a different anti-Ring1b antibody. As expected, we found that Ring1b is similarly bound to target promoters in G1 and G2 phases (Figure S1K), and that the antibody specifically recognises Ring1b, because the binding signal is lost in mESCs that are genetic null for *Ring1b* (Figure S1L, M, N). Although sustained recruitment of Ring1b to target promoters across interphase might suggest continued repression of Ring1b target genes in G1, S and G2 phases, analysis of nascent RNA datasets, obtained by 4-thiouridine labelling followed by sequencing (4sU-seq) ^19^, revealed that Ring1b target genes are transcriptionally downregulated during S and G2 phases compared to G1 phase (Figure 1F). Augmented RNA synthesis in G1 compared to G2 phase was particularly evident at lowly expressed bivalent genes (Figure 1G, H, I). We concluded that although Ring1b remains bound to promoter regions during interphase, its target genes display enhanced transcriptional repression specifically during S and G2 phases.

**Figure 1.**
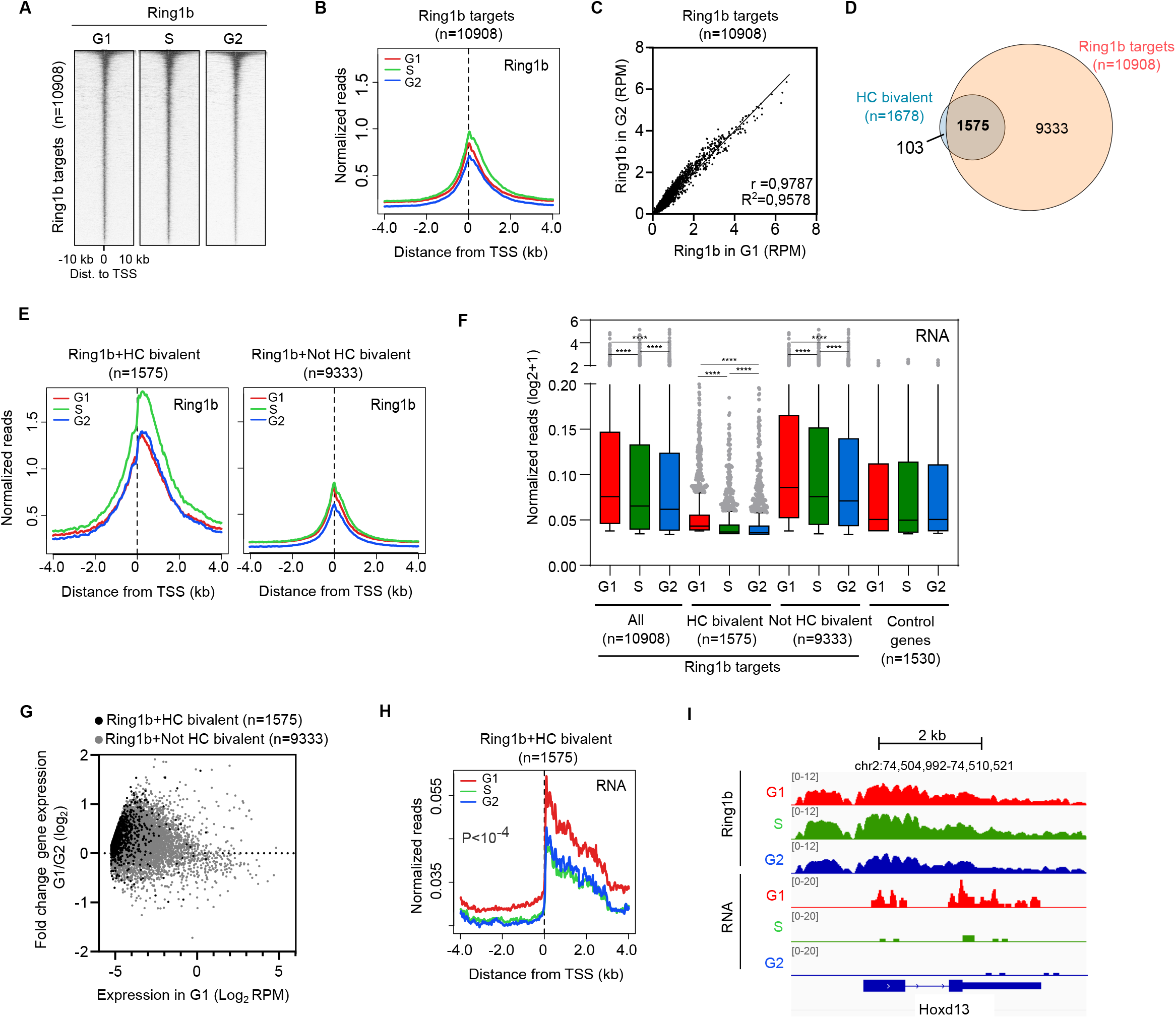
Ring1b remains bound to target gene promoters across interphase. **(A)** Heatmaps of normalized Ring1b ChIP-seq reads around the TSS (±10 kb) of target promoters (n=10908) at different phases of the cell cycle. Genes are ranked according to the signal in G2 phase. **(B)** Average binding profile of Ring1b around the TSS of Ring1b target promoters in G1 (red), S (green) and G2 (blue) phases. **(C)** Correlation analysis between the signal of Ring1b binding around the TSS (−0.5 kb to +1.5 kb) of target genes during G1 (x axis) and G2 (y axis). Linear regression R^2^ and pearson coefficient (r) are indicated. **(D)** Venn diagram showing the overlap between HC bivalent promoters (blue) and Ring1b target promoters (orange). HC bivalent promoters were previously defined ^19^. **(E)** Average binding profile of Ring1b around the TSS of target genes that are HC bivalent promoters (left panel) or not (right panel) defined in figure 1D in G1 (red), S (green) and G2 (blue) phases. **(F)** Boxplots comparing 4sU-seq RNA reads (RPM) mapped to the proximal promoter region (TSS to +3Kb) of Ring1b targets depending on whether they are HC bivalent or not in the indicated cell cycle phases. Genes that are not bound by Ring1b (0.2 > Log_2_ FC G1/G2 > -0.2) are shown as a control. Asterisks (*) mark statistically significant differences. **(G)** MA plot of fold change gene expression between cells in G1 and G2 phases (4sU-seq normalized reads mapping from the TSS to +3kb) at HC bivalent Ring1b targets (black dots) and Not HC bivalent Ring1b targets (grey dots). Nascent RNA expression in G1 is represented in the x-axis. **(H)** Average 4sU-seq RNA reads at Ring1b+HC bivalent promoters in G1 (red), S (green) and G2 (blue) phases. *P* indicates ANOVA test p-value. **(I)** Genome browser view of Ring1b binding and nascent RNA synthesis at indicated cell cycle phases at the bivalent gene *Hoxd13*.

### Increased recruitment of vPRC1 to target genes during S and G2 phases is associated with accumulation of H2AK119ub1 and enhanced transcriptional repression

To study whether PRC1 is responsible for enhanced gene repression upon G1 exit at Ring1b target genes, we focused on the analysis of vPRC1, that harbour most of the transcriptional repression capacity of the polycomb system through H2AK119ub1 ^33-35^. In mESCs, vPRC1 complexes are characterized by the presence of Rybp ^33, 36^(Figure 2A). Analysis of the genome-wide distribution of Rybp revealed that, in stark contrast to Ring1b, recruitment of Rybp to target regions was markedly increased during S and G2 phases as compared to G1 (Figure S2A, B). Rybp was mostly bound to gene promoter regions (n=10925) (Figure S2A, C) where its recruitment around the TSS was clearly increased upon G1 exit (Figure 2B). In agreement with previous reports ^36, 37^, most of Rybp targets (9122 out of 10925 genes) were also bound by Ring1b (Ring1b+Rybp targets) (Figure 2C). Although recruitment of Rybp to Ring1b+Rybp target promoters was enhanced during interphase progression, Ring1b remained bound at similar levels at all phases of the cell cycle (Figure 2D, S2D). Importantly, ChIP-seq analysis of H2AK119ub1 demonstrated that enhanced recruitment of Rybp during S and G2 phases is associated to accumulation of H2AK119ub1 at Ring1b+Rybp target genes (Figure 2E, F, G,

**Figure 2.**
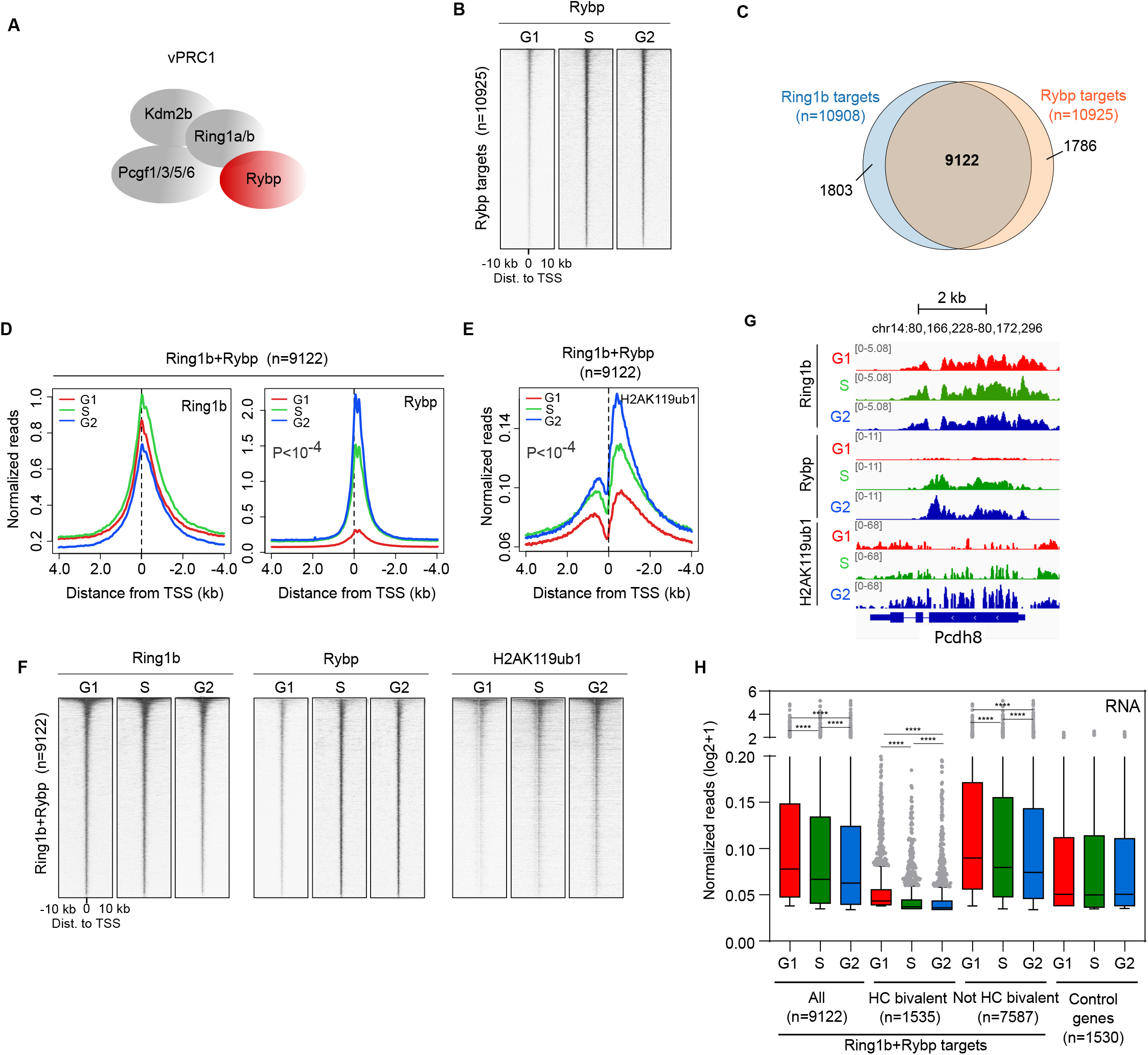
Increased recruitment of Rybp to target genes during S and G2 phases is associated with accumulation of H2AK119ub1 and enhanced transcriptional repression. **(A)** Scheme highlighting Rybp (red) in vPRC1 complexes in mESCs. **(B)** Heatmaps of normalized Rybp ChIP-seq reads around the TSS (±10 kb) of target promoters (n=10925) at different phases of the cell cycle. Genes are ranked according to the signal in G2 phase. **(C)** Venn diagram showing the overlap between Ring1b (blue) and Rybp (orange) target promoters. **(D, E)** Average binding profile of Ring1b and Rybp (D) and H2AK119ub1 (E) around the TSS of promoters bound by Ring1b and Rybp (identified in figure 2C) in G1 (red), S (green) and G2 (blue) phases. *P* indicates ANOVA test p-value. (**F**)Heatmaps of normalized Ring1b, Rybp and H2AK119ub1 ChIP-seq reads around the TSS (±10 kb) of Ring1b and Rybp target promoters at different phases of the cell cycle. Genes are ranked according to the signal of Ring1b in G2 phase. **(G**)Genome browser view of Ring1b, Rybp and H2AK119ub1 enrichment at the target gene *Pcdh8* during G1, S and G2 phases. **(H**) Boxplots comparing 4sU-seq RNA reads (RPM) mapped to the proximal promoter region (TSS to +3Kb) of Ring1b and Rybp targets depending on whether they are HC bivalent or not (as defined in figure S2G) in the indicated cell cycle phases. Genes that are not bound by Ring1b+Rybp (0.2 > Log_2_ FC G1/G2 > -0.2) are shown as a control. Asterisks (*) mark statistically significant differences.

S2E). Although the binding of Rybp and H2AK119ub1 in G1 phase is very low as compared to the G2 phase (Figure 2D), it is still noticeable when compared to promoter regions that are not PRC1-targets and are heavily methylated in their DNA (Figure S2F). Thus, low levels of Rybp and H2AK119ub1 present in G1 phase might be enough to trigger a positive feedback recruitment mechanism (i.e., through Rybp) that is time dependent and leads to the accumulation of vPRC1 complexes during S and G2. The overlap between genes bound by Ring1b and Rybp is very high (Figure 2C) and therefore we expectedly found that, similarly to Ring1b target promoters, Ring1b+Rybp genes are transcriptionally repressed upon G1 exit, and that this effect is particularly obvious at lowly expressed bivalent genes (Figure 2H, S2G, H, I). Taken together, these results indicate that vPRC1 is accumulated at target promoters during S and G2 phases, and that this is coupled to accumulation of H2AK119ub1 and enhanced transcriptional repression.

### Accumulation of cPRC1 at target genes is coupled to strengthened promoter-promoter 3D contacts during G2 phase

In mESCs, cPRC1 contain Phc1 and Cbx7 subunits ^38, 39^, facilitating recruitment of PRC1 to H3K27me3-target genes ^40, 41^ and mediating DNA interactions among polycomb-bound genes ^42, 43^ (Figure 3A). To ask whether recruitment of cPRC1 is also enhanced during S and G2 phases we analysed the genome-wide distribution of Cbx7 by ChIP-seq in cell cycle sorted mESCs. Recruitment of Cbx7 was clearly augmented in cells in G2 as compared to cells in G1 phase (Figure S3A, B). Cbx7 was preferentially bound to gene promoter regions (n=2395), and these were highly coincident at the three different phases of interphase (Figure S3C). In fitting with previous studies ^38^, most Cbx7-bound promoters (2340 out of 2395) were also targeted by Ring1b, but many Ring1b targets were not bound by Cbx7, indicating that cPRC1 is only present in a subset of genes targeted by vPRC1 (Figure 3B). Ring1b+Cbx7 target promoters (n=2340) showed a remarkable increase in Cbx7 binding in G2 phase as compared to G1 phase (Figure 3C, D, S3D). Recruitment of Ring1b at both phases of the cell cycle remained constant but it was higher at Cbx7-bound gene promoters compared to non-bound ones (compare Ring1b+Cbx7 to Ring1b only panels in figure 3D, S3D), suggesting that Cbx7 facilitates the accumulation of Ring1b at target promoters. We concluded that recruitment of Cbx7 to target promoters is enhanced in G2 phase compared to G1 phase.

**Figure 3.**
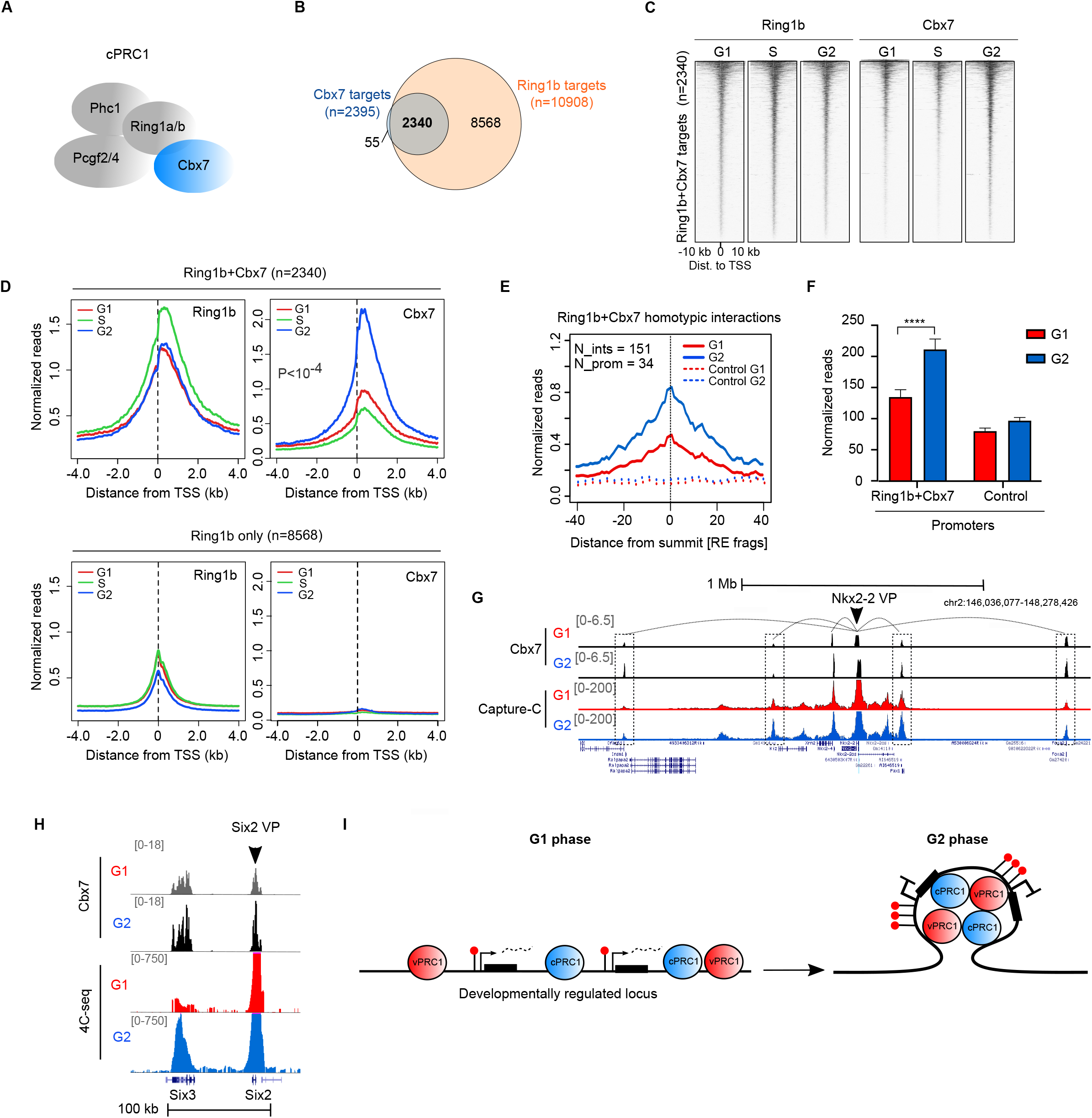
Accumulation of Cbx7 at target genes is coupled to strengthen promoter-promoter contacts during G2 phase. **(A)** Scheme highlighting Cbx7 (blue) in cPRC1 complexes in mESCs. **(B)** Venn diagram showing the overlap between Ring1b (orange) and Cbx7 (blue) target promoters. **(C)** Heatmaps of normalized Ring1b and Cbx7 ChIP-seq reads around the TSS (±10 kb) of promoters bound by Ring1b and Cbx7 (as identified in figure 3B) at different cell cycle phases. Genes are ranked according to the signal of Ring1b in G2 phase. **(D)** Average binding profile of Ring1b and Cbx7 around the TSS of Ring1b-bound promoters that are targeted (top panel) or not (bottom panel) by Cbx7 in G1 (red), S (green) and G2 (blue) phases. *P* indicates ANOVA test p-value **(E)** Average normalized reads of 3D chromatin interactions (number of interactions = 151) measured by capture-C of Ring1b+Cbx7-bound regions involving at least one promoter (number of promoters=34) in G1 (red) and G2 (blue) phases. The X axis represents the distance from the summit of the interaction peak measured as a function of DpnII restriction fragments. Dashed lines show enrichment at distance-matched control sites from each promoter and interaction in the opposite direction. **(F)** Capture-C average normalized reads of interactions involving Ring1b+Cbx7 promoters (promoters = 34; interactions = 151) compared to a group of control promoters not bound by Ring1b+Cbx7 (promoters = 41; interactions = 261) in G1 (red) and G2 (blue) phases. Asterisks (*) mark statistically significant differences. **(G)** Genome browser view of Capture-C interactions involving the Nkx2-2 promoter (viewpoint, VP) in G1 (red) and G2 (blue) phases. Binding of Cbx7 in G1 and G2 phases is shown. Dashed boxes highlight most obvious interacting chromatin regions. **(H)** 4C-seq analysis of the interaction between the *Six2* (viewpoint, VP) and *Six3* promoters in G1 (red) and G2 (blue) phases. Binding of Cbx7 in G1 and G2 phases is shown. **(I)** Schematic diagram of observations described in this manuscript. Black boxes indicate genes, dashed lines, nascent RNA and red lollipops H2AK119ub1. PRC1 complexes are represented as blue or red spheres. PRC1 complexes accumulate around target promoters in G2 compared to G1 phase, leading to enhanced ubiquitination of H2AK119, stronger 3D interactions and firmer gene repression.

To address whether changes in Cbx7 recruitment during cell cycle transition are associated to topological changes of the genome, we compared 3D interactions of polycomb target genes in cells in G1 and G2 phases using capture-C. Gene promoters bound by Ring1b+Cbx7 displayed strengthened interactions with other DNA regions in cells in G2 phase compared to cells in G1 phase (Figure S3E). Remarkably, increased 3D interactions in G2 were evident among loci that were bound by Ring1b+Cbx7 (homotypic interactions) (Figure 3E, F). Changes in interactions in G2 phase were appreciable at individual loci (i.e. *Nkx2-2* gene, Figure 3G, see regions highlighted by dotted lines), and were confirmed using 4C-seq at the *Six2/Six3* locus (Figure 3H). Although interactions of cPRC1-bound regions were strengthened in G2 phase, they were already evident in G1 phase when compared to negative control regions (Figure 3E). This indicates that 3D interactions mediated by cPRC1 are present in G1, but that they are intensified as cells transit into G2 phase. In fitting, Cbx7 binds to target promoters in G1 phase (Figure S3F), but its recruitment is augmented in G2 phase (Figure 3D). Taken together, our analyses indicate that both vPRC1 and cPRC1 complexes, as well as their associated functional effects - H2AK119ub1, gene repression and 3D chromatin interactions - are enhanced upon G1 exit in mESCs (Figure 3I).

### Developmentally regulated transcription factors are common targets of PRC2/cPRC1/vPRC1 that display enhanced recruitment of PRC1 during G2 compared to G1 phase

Our analyses indicate that most promoters targeted by Ring1b can also recruit Rybp but that only a proportion of them are bound by Cbx7 (Figure 2C and 3B). In agreement with previous studies ^37, 44^, comparative analysis of PRC target regions revealed two major groups of PRC1 target promoters: 2093 promoters that were bound by both PRC1 and PRC2 (vPRC1/cPRC1/PRC2 targets), and 6222 genes that were bound by vPRC1 complexes only (vPRC1 specific targets). vPRC1/cPRC1/PRC2 targets were enriched in developmentally regulated transcription factors, while the vPRC1-specific ones were enriched in metabolism and signalling processes (Figure 4A). Recruitment of Ring1b, Rybp, Cbx7 and H2AK119ub1 to promoter regions was higher in vPRC1/cPRC1/PRC2 targets compared to vPRC1-specific ones (Figure 4B, S4A, S4B, S4C), probably reflecting positive feedback mechanisms that facilitate accumulation of PRCs and formation of more extensive repressive domains at this subset of vPRC1-targeted genomic regions. In agreement, vPRC1/cPRC1/PRC2 targets were expressed at a very low level compared to vPRC1-specific genes (Figure 4C). Importantly, the extent to which recruitment of PRC1 was enhanced in G2 phase compared to G1 was higher in vPRC1/cPRC1/PRC2 target promoters than in vPRC1-specific ones (Figure 4D), indicating that the cell-cycle-dependent regulation of PRC1 is particularly effective at chromatin domains that are highly enriched for PRC1 binding. Interestingly, changes in RNA synthesis were also more homogeneous at vPRC1/cPRC1/PRC2 targets than at vPRC1-specific genes (Figure 4E), supporting that high levels of PRC1 are linked to more robust transcriptional repression of target genes in G2 phase. We concluded that developmentally regulated transcription factors display the most obvious cell-cycle-dependent regulation of vPRC1, cPRC1 and PRC2 binding, and that this is coupled to enhanced transcriptional repression during G2, as compared to the G1 phase.

**Figure 4.**
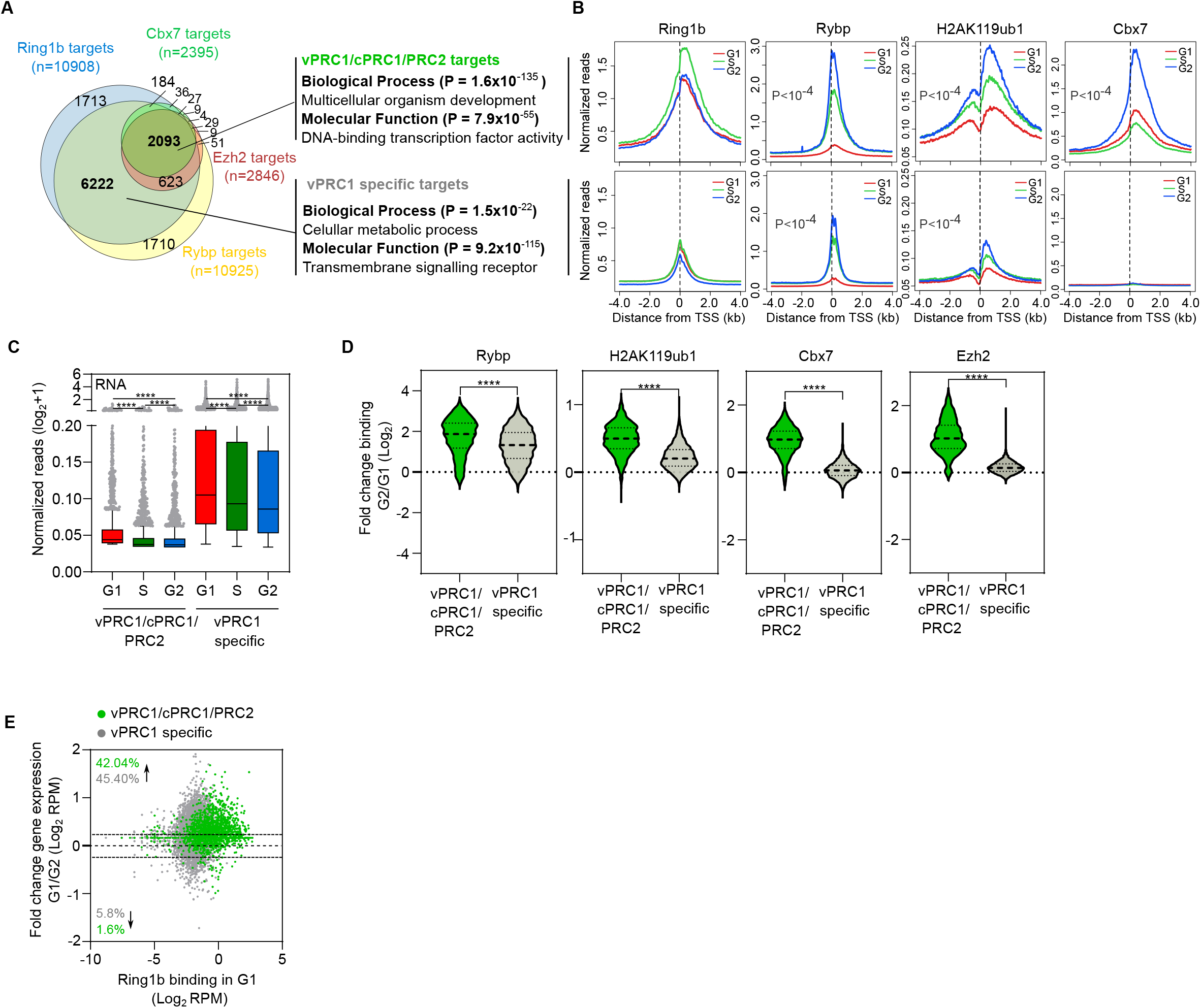
Developmentally regulated transcription factors are common targets of PRC2/cPRC1/vPRC1 that display enhanced recruitment of PRC1 during G2 compared to the G1 phase. **(A)** Venn diagram showing the overlap between Ring1b (blue), Rybp (yellow), Cbx7 (green) and Ezh2 (red) target promoters. Gene Ontology analyses of gene promoters bound by vPRC1, cPRC1 and PRC2 (n=2093) or by vPRC1 only (n=6222) are shown on the right. **(B)** Average binding profile around the TSS of Ring1b, Rybp, H2AK119ub1 and Cbx7 at vPRC1/cPRC1/PRC2 (top row) or vPRC1 specific (bottom row) target genes (as defined in figure 4A) in G1 (red), S (green) and G2 (blue) phases. *P* indicates ANOVA test p-value. **(C)**Boxplots comparing 4sU-seq nascent RNA reads (RPMs, TSS to +3Kb) of vPRC1/cPRC1/PRC2 targets (n=2093) and vPRC1 specific targets (n=6222) at indicated phases of the cell cycle. Asterisks (*) mark statistically significant differences. **(D)**Violin plots showing the fold change binding between G2 and G1 of Rybp, H2AK119ub1, Cbx7 and Ezh2 at promoter regions (−0.5 kb to +1.5 kb relative to TSS) of vPRC1/cPRC1/PRC2 (green) and vPRC1 specific target genes (grey). Asterisks (*) mark statistically significant differences. **(E)**MA plot showing fold change gene expression between cells in G1 and G2 phases (4sU-seq normalized reads mapping from the TSS to +3kb) at vPRC1/cPRC1/PRC2 (green dots) and vPRC1 specific target promoters (grey dots). Binding of Ring1b in G1 is represented in the x-axis. Percentage of up- or down-regulated (above threshold FC>1.2) genes for are indicated.

### Depletion of Ring1b perturbs 3D chromatin interactions and transcriptional repression in G2 phase

To address whether the accumulation of PRC1 drives strengthened chromatin interactions and enhanced gene repression during G2 phase, we analysed whether depletion of Ring1b protein perturbs these features at PRC1-bound genes at different phases of the cell cycle. We introduced the FUCCI reporter system into previously derived *Ring1b* conditional knockout mESCs (*Ring1a*^−/–^ ;*Ring1b*^fl/fl^;*Rosa26*::*CreERT2)* (FUCCI-*Ring1b*^fl/fl^) ^45^. As expected, treatment with tamoxifen (Tmx) led to undetectable levels of Ring1b protein and H2AK119ub1 within seventy-two hours in the new FUCCI expressing genetic clone (Figure S1L, S1M, S1N). Parental control (FUCCI-*Ring1b*^fl/fl^) and tamoxifen-treated cells (FUCCI-*Ring1b*^-/-^) were sorted by flow cytometry and populations enriched for cells in G1 or G2 phases were subjected to capture-C analyses (Figure 5A). Homotypic interactions of cPRC1 binding sites were severely reduced to background levels in FUCCI-*Ring1b*^-/-^ cells in either G1 or G2 phases (Figure 5B), while interactions of control regions were not affected by the depletion of Ring1b (Figure S5A). Loss of 3D interactions upon Ring1b depletion was also evident at individual loci (i.e. *Nkx2-2* gene, Figure 5C, see regions highlighted by dashed lines). Consequently, we established that Ring1b is required to maintain the topological organization of polycomb-bound regions in G1 and G2 phases.

**Figure 5.**
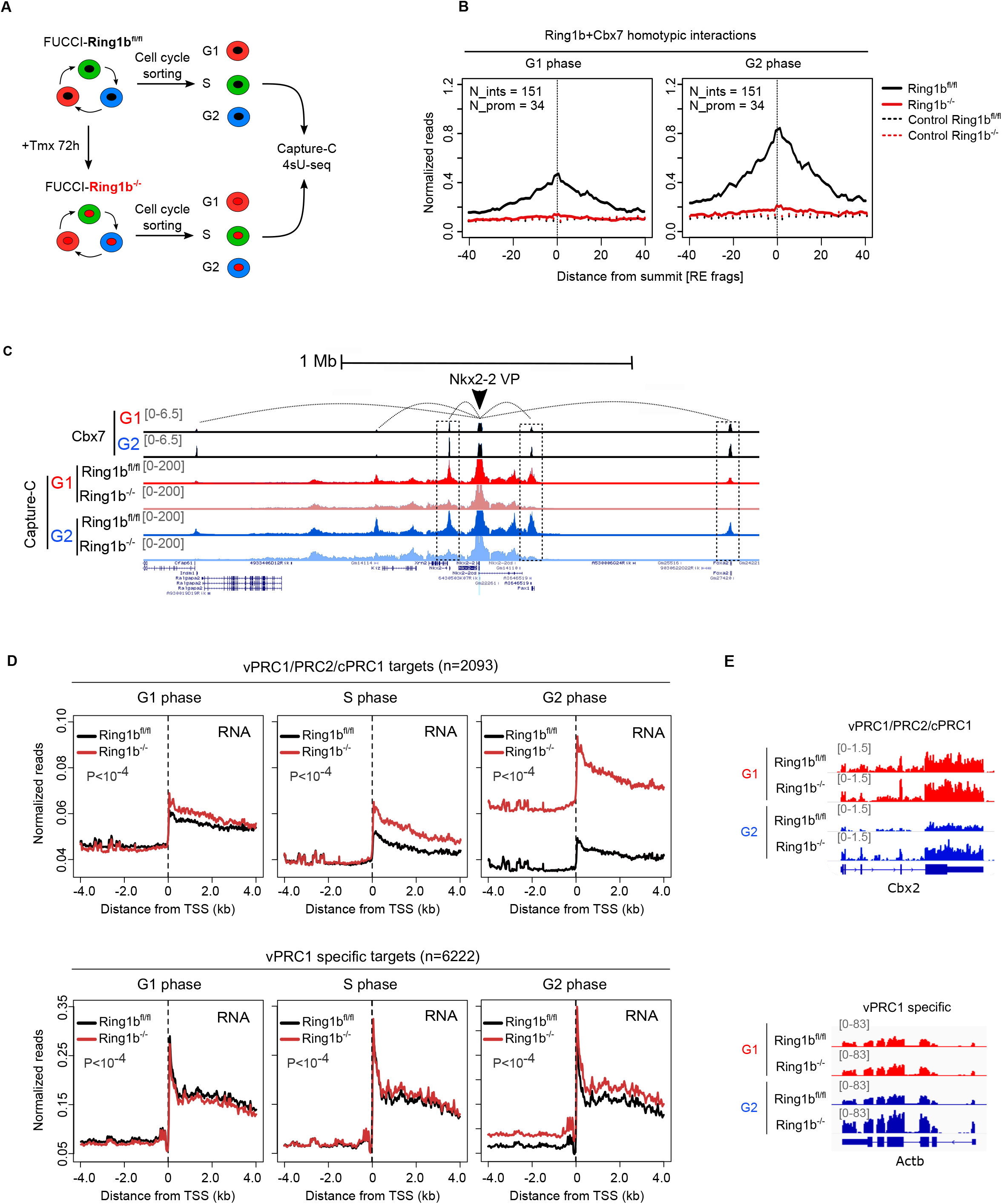
Depletion of Ring1b perturbs 3D chromatin interactions and transcriptional repression in G2 phase. **(A)**Scheme representing the experimental setup to study 3D chromatin interactions and nascent RNA in PRC1 null cells. Ring1a^-/-^; Ring1b^fl/fl^; Rosa26::CreERT2;FUCCI (FUCCI-Ring1b^fl/fl^) mESCs were treated with Tmx for 72h, cell cycle sorted and subjected to Capture-C and 4sU-seq analyses. **(B)**Average normalized reads of 3D chromatin interactions (number of interactions = 151) measured by Capture-C of Ring1b+Cbx7-bound regions involving at least one promoter (number of promoters=34) in Ring1b^fl/fl^ (black line) and Ring1b^-/-^ (red line) cells during G1 (left) and G2 (right) phases. The X axis represents the distance from the summit of the interaction peak measured as a function of DpnII restriction fragments. Dashed lines show enrichment at distance-matched control sites from each promoter and interaction in the opposite direction. **(C)**Genome browser view of Capture-C interactions involving the Nkx2-2 promoter (viewpoint, VP) in Ring1b^fl/fl^ and Ring1b^-/-^ cells during G1 (red) and G2 (blue) phases. Binding of Cbx7 in G1 and G2 phases is shown. Dashed boxes highlight interactions that are lost in Ring1b^-/-^ compared to the Ring1b^fl/fl^ cells. **(D)**Average nascent RNA reads at vPRC1/cPRC1/PRC2 (top panel) and vPRC1 specific (bottom panel) target promoters in G1 (left), S (middle) and G2 (right) phases in *Ring1b*^fl/fl^ (black lines) and *Ring1b*^-/-^ (red lines) mESCs. P indicates ANOVA test p-value **(E)**Genome browser view of nascent RNA at indicated cell cycle phases in Ring1b^fl/fl^ and Ring1b^-/-^ mESCs at the vPRC1/cPRC1/PRC2 target gene *Cbx2* (top panel) and the vPRC1 specific target gene *Actb* (bottom panel).

Next, we asked whether Ring1b is required to maintain transcriptional repression of PRC1-target genes at the different phases of interphase. Upon tamoxifen treatment and cell cycle sorting, we measure nascent RNA synthesis using 4sU-seq in FUCCI-*Ring1b*^-/-^ and parental control cells in G1, S and G2 phases (Figure 5A). Depletion of Ring1b protein led to a drastic upregulation of most vPRC1/cPRC1/PRC2 targets in G2 phase (1898 genes out of 2093 were transcriptionally derepressed) (Figure 5D, S5B, S5C). The loss of Ring1b also affected the regulation of vPRC1-specific genes in G2 phase (2139 genes out of 6222 were upregulated) (Figure 5D, S5B, S5C), albeit the number of affected genes and the magnitude of mis-regulation was dimmed compared to vPRC1/cPRC1/PRC2 targets (Figure S5D, S5E). Strikingly, depletion of Ring1b barely perturbed the transcriptional repression of its target genes in G1 phase, while cells in S phase displayed and intermediate phenotype (Figure 5D, S5C). The effect of Ring1b depletion in nascent RNA synthesis was evident at individual loci in G2 cells (Figure 5E). Therefore, we concluded that that Ring1b repressive activity builds up during S and G2 phases in coordination with augmented binding of Rybp and H2AK119ub1 to Ring1b target genes (Figure 2D, E). Nevertheless, we cannot fully discard that de-repression of Ring1b target genes during S and G2 phases is partly a consequence of the action of transcriptional activation signals specifically present during these phases.

### Ring1b regulates the cell-cycle-dependent induction of gene transcription during lineage transition in mESCs

We hypothesized that reduced transcriptional repression of developmental regulators by PRC1 in the G1 phase could facilitate their activation in response to differentiation signals. To test this, we used published a mESCs line that express endogenous levels of functional Ring1b proteins fused to the auxin-inducible degron (AID::Ring1b) that can be conditionally and rapidly degraded upon two hours of treatment with auxin (IAA) ^46^. We introduced the FUCCI plasmid in these cells to establish a system (AID::Ring1b;FUCCI) in which we could deplete Ring1b in cells at a specific phase of the cell cycle and analyse their response to retinoic acid (RA) stimulation. AID::Ring1b;FUCCI mESCs were sorted in G1, S and G2 phases and plated in differentiation conditions (without LIF and in the presence of retinoic acid (RA)), in the presence (IAA+, Ring1b-depleted) or in the absence (UNT, control) of IAA (Figure 6A, B). Cell cycle sorted cells were collected six hours after plating, when most of the cells remained at the same phase of the cell cycle at which they were initially plated (Figure S6A). Then, we performed mRNA-seq to analyse how the depletion of Ring1b at specific phases of the cell cycle affected the transcriptional induction of target genes early during lineage transition. Principal component analysis readily discriminated samples depending on RA stimulation, Ring1b depletion and the phase of the cell cycle (figure S6B), demonstrating that these three variables regulate gene transcription in this system. Treatment of cells with RA during six hours induced the expression of 881 genes that were associated to the three different germ layers, with a preference towards the ectoderm lineage (Figure S6C, S6D), indicating that cells are exiting pluripotency and transiting into early lineage specification. In fitting with the repressive role of PRC1, depletion of Ring1b prompted upregulation of more than 400 genes in IAA-treated compared to untreated cells, most of which displayed high levels of Ring1b and enrichment of H2AK119ub1 at their promoter regions (Figure S6E). No gene was significantly downregulated in Ring1b-depleted cells compared to the untreated control (Figure S6E), demonstrating that in this context, Ring1b functions as a transcriptional repressor. Clustering of differentially expressed genes in control and IAA-treated cells showed that genes de-repressed upon IAA treatment (cluster II) were highly enriched in Ring1b, Rybp, Cbx7 and H2AK119ub1 as compared to genes whose gene expression profile did not change upon depletion of Ring1b (cluster I and III), indicating that cluster II is highly enriched for direct targets of Ring1b (Figure S6F, S6G). Importantly, a large subset of Ring1b-target genes displayed higher expression in G1 relative to S and G2 phases in parental untreated cells after stimulation with RA (371 genes identified in Cluster II-A, right panel in Figure S6F), indicating that Ring1b regulates the cell-cycle-dependent activation of genes upon retinoic acid stimulation.

**Figure 6.**
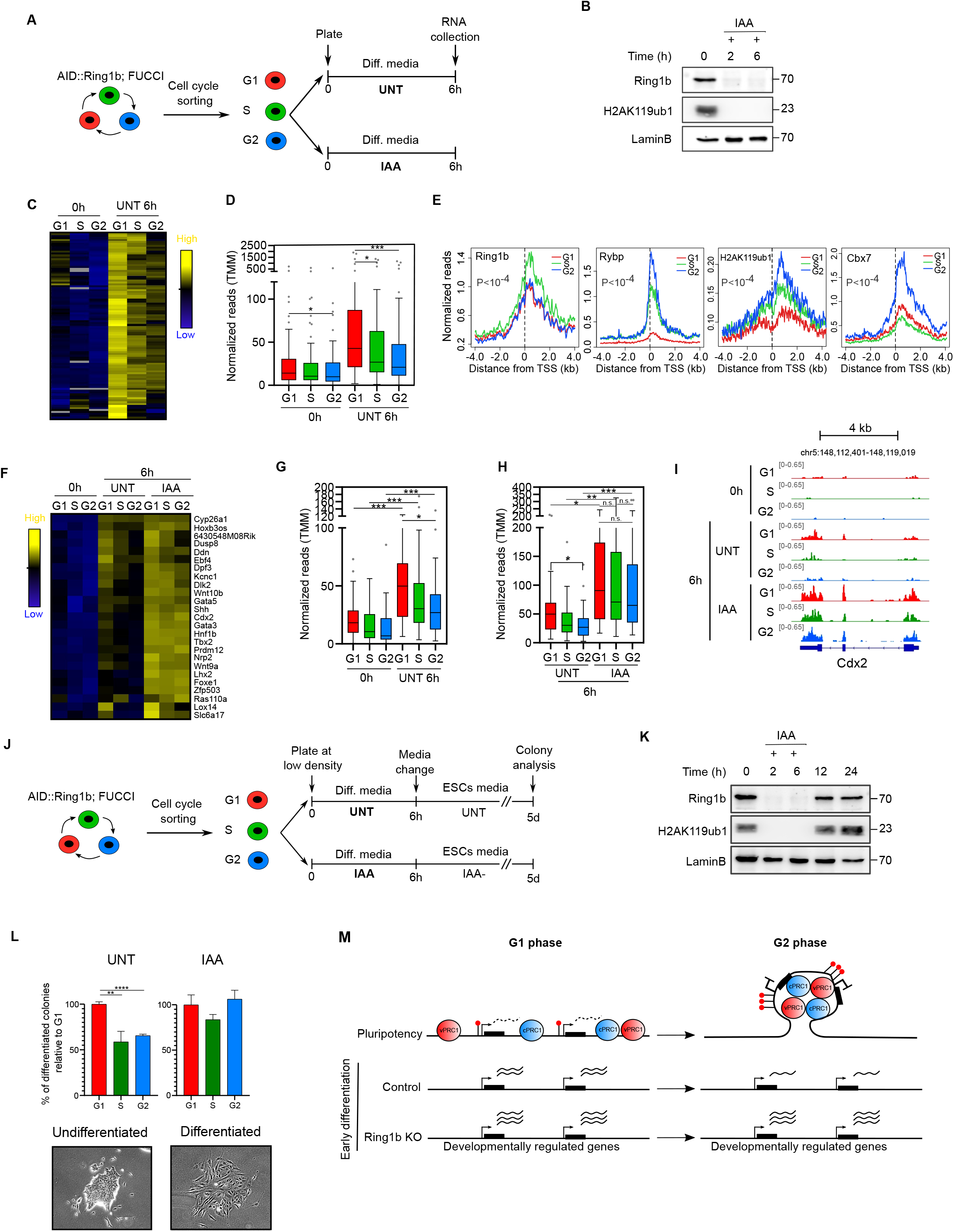
Ring1b regulates the cell-cycle-dependent induction of gene transcription during lineage transition in mESCs. **(A)**Scheme representing the experimental strategy to analyse the cell cycle phase-specific function of Ring1b during lineage. AID::Ring1b;FUCCI mESCs were cell cycle sorted and plated in differentiation media (5 µM RA in the absence of LIF) with (IAA+) or without (control UNT) auxin (IAA) during 6h before RNA collection. **(B)**Western blots of whole cell extracts for Ring1b and H2AK119ub1 proteins in AID::Ring1b;FUCCI mESCs treated with IAA for the indicated times. Lamin B was used as loading control. Molecular weight is indicated in kDa. **(C)**Hierarchical clustering of mRNA expression levels (TMM) of cell cycle sorted samples at 0h and 6h after RA treatment of genes preferentially upregulated during G1 compared to G2 phase (FC>2, n=97). **(C)**Boxplot of mRNA expression levels (TMM) of cell cycle sorted samples at 0h and 6h after RA treatment of genes preferentially upregulated during G1 compared to G2 phase (n=97). Asterisks (*) mark statistically significant differences. **(D)**Average binding profile of Ring1b, Rybp, H2AK119ub1 and Cbx7 around the TSS of genes preferentially upregulated during G1 compared to G2 (n=97) in G1, S and G2 phases. *P* indicates ANOVA test p-value. **(E)**Hierarchical clustering of mRNA expression (TMM) of cell cycle sorted samples at 0h and 6h after RA treatment in Ring1b-depleted and control UNT cells. Genes preferentially upregulated during G1 compared to G2 phase that display higher levels of H2AK119ub1 (identified in figure S6G, n=25) are shown. Gene symbols are indicated. **(F)**Boxplot of mRNA expression (TMM) at 0h and 6h after RA treatment in cell cycle sorted cells. Genes preferentially upregulated during G1 compared to G2 phase that display higher levels of H2AK119ub1 (identified in figure S6G, n=25) are shown. Asterisks (*) mark statistically significant differences. **(G)**Boxplot of mRNA expression (TMM) at 6h after RA treatment in cell cycle sorted Ring1b-depleted and control UNT cells. Genes preferentially upregulated during G1 compared to G2 phase that display higher levels of H2AK119ub1 (identified in figure S6G, n=25) are shown. Asterisks (*) or n.s. mark statistically significant or not significant differences respectively. **(H)**Genome browser view of mRNA expression at indicated phases of the cell cycle at 0h and 6h after RA stimulation in cell cycle sorted Ring1b-depleted and control UNT cells at the *Cdx2*. **(I)**Scheme showing the experimental design to study the cell cycle phase-specific role of Ring1b in productive cell differentiation. AID::Ring1b;FUCCI mESCs were cell cycle sorted and plated at low density in differentiation media (0.1 µM RA in the absence of LIF) with (Ring1b-depleted) or without (control UNT) auxin (IAA). After 6h cells were cultured in standard ESCs media and colonies were fixed and analysed after five days. **(J)**Western blots of whole cell extracts for Ring1b and H2AK119ub1 proteins in AID::Ring1b;FUCCI mESCs treated with IAA for the indicated times. Lamin B was used as loading control. Molecular weight is indicated in kDa. **(K)**Histograms showing the percentage of differentiated colonies relative to G1 obtained after five days from cell cycle sorted populations in parental control (UNT) and IAA-treated (Ring1b-depleted) cells (see experimental setup in figure 6J). Pictures showing examples of colonies with undifferentiated (compact and 3D colonies with round cells) or differentiated (flat and spread colonies with elongated cells) morphologies are shown. Asterisks (*) mark statistically significant differences. **(L)**Scheme summarizing the observations made in this manuscript. Black boxes indicate genes, red lollipops H2AK119ub1 and PRC1 complexes are represented as blue or red spheres. RNA expression is represented as dashed (leaky expression) or solid (higher expression) wavy lines. PRC1 complexes accumulate around target promoters in G2 compared to G1 phase, leading to enhanced ubiquitination of H2AK119, stronger 3D interactions and firmer gene repression. Upon induction of cell differentiation PRC1 target genes are preferentially activated during G1 phase. This cell cycle phase-dependent activation is perturbed in Ring1b-depleted cells.

To analyse the role of Ring1b in the regulation of cell-cycle-phase-dependent transcriptional activation of differentiation genes we identified ninety-seven genes that displayed solid preferential transcriptional activation in G1 relative to G2 phase in untreated cells (FC>2, FDR<0.05) (Figure 6C, 6D). Promoter of these genes displayed expected binding by cPRC1 and vPRC1 around their TSS in pluripotent cells, and displayed enhanced recruitment of Cbx7 and Rybp proteins in G2 phase compared to G1 phase (Figure 6E). Likewise, these genes displayed accumulation of H2AK119ub1 at their promoter regions in S and G2 phases (Figure 6E). Because genes with high level of Ring1b and H2AK119ub1 were the more responsive to Ring1b deletion (Figure S6E), we focused our analysis on the subset of cell-cycle-regulated genes that displayed higher levels of H2AK119ub1 (Q1 in Figure S6H). These included well-known DNA binding proteins involved in cell differentiation such as Ddn, Ebf4, Dpf3, Gata5, Cdx2, Gata3, Hnf1b, Tbx2, Lhx2, Foxe1 and Zfp503. As expected, stimulation with RA induced higher expression of these genes in G1 compared to cells in S and G2 phases in control cells (Figure 6F, 6G). Remarkably, cells depleted for Ring1b showed similarly abnormally high levels of RNA transcripts irrespectively of the phases of the cell cycle in which they were induced with RA (Figure 6F, 6H, 6I). Thus, the increased transcriptional activation of Ring1b-target genes of G1 parental cells in response to RA stimulation was mostly lost in Ring1b-depleted cells, supporting that Ring1b is required to dim the transcriptional induction of target genes in response to RA during S and G2 phases. In fitting, this was traduced in more accused de-repression of target genes in Ring1b-depleted cells, compared to UNT cells, in S and G2 phases than in G1 phase (Figure S6I). We concluded that, upon receival of the RA differentiation stimulus, mESCs require PRC1 to hamper the transcriptional activation of target genes during S and G2 phases.

To test whether differences observed during early lineage transition translate into changes in effective cell differentiation, we plated cells at low density in differentiation media upon cytometry sorting in G1, S and G2 phases, we stimulated them with RA and switched them back to grow in standard ESCs media during five more days (Figure 6J). Colonies were then classified as undifferentiated or differentiated under the microscope (Figure 6J). This experimental setup allowed us to analyse the extent to which transient stimulation with RA induced irreversible cell differentiation in cells in G1, S or G2 phases. Importantly, wash out of IAA from the culture media upon IAA-treatment resulted in rapid restoration of Ring1b and H2AK119ub1 levels (Figure 6K). Therefore, by co-treating cell cycle sorted AID::Ring1b;FUCCI cells with IAA and RA during the initial six hours we could analyse whether depletion of Ring1b affected the cell-cycle-phase-dependent response of pluripotent cells to RA stimulation (Figure 6J). In agreement with previous analyses ^14^, control cells formed substantially more fully differentiated colonies in G1 compared to cells in S and G2 phases (Figure 6L). Remarkably, cells that were depleted of Ring1b specifically during RA stimulation gave rise to differentiated colonies with a similar efficacy, independently of the phase of the cell cycle at which they received the differentiation stimulus (Figure 6L). Therefore, we concluded that PRC1 activity is required to maintain the ability of mESCs to trigger effective cell differentiation preferentially in G1 phase (Figure 6M).

## Discussion

Our study reveals that PRC1 repressive activity is enhanced in S and G2 phases compared to the G1 phase in pluripotent cells. This is enabled by the accumulation of Rybp and H2AK119ub1 at target promoters which leads to firmer transcriptional repression. In addition, we found that the PRC1 catalytic subunit Ring1b is required to facilitate the cell-cycle-phase-specific activation of differentiation genes in response to RA stimulation in mESCs. Hence, our results strongly support that incremental enrolment of PRC1 activity during S and G2 reduces the tendency of cells to respond to developmental cues, facilitating the transcriptional induction of cell differentiation genes in the G1 phase. These discoveries are noteworthy for two reasons. Because we reveal that cell cycle regulation of PRC1 activity is a hitherto unrecognized key functional aspect of the polycomb repressive system in mammals, and because we disclose for the first time the molecular mechanism by which an epigenetic factor regulates the cell-cycle-phase-specific cell differentiation capacity of stem cells.

Repressive histone post-translational modifications (PTMs), H3K9me3 and H3K27me3, have been proposed to self-propagate through DNA synthesis using a read-write mechanism ^47-49^. In agreement with this model, we found that the total amount of H2AK119ub1 on chromatin was similar during G1, S and G2 phases. Notwithstanding, we found that there are major changes in the genome-wide distribution of H2AK119ub1 during interphase progression. Thus, our data imply that most of H2AK119ub1 is not bound to gene regulatory elements but to other genomic regions, in fitting with the pervasive low levels of H2AK119ub1 detected around the genome ^35^. Importantly, the distribution of PRC1 proteins, rather than their global amount, is perhaps the critical regulatory factor, because polycomb function largely rely on the accumulation or PRCs at target genomic regions to form repressive chromatin domains ^20, 21^.

Polycomb domains formation is based on positive feedback loops that involve crosstalk between different PRCs. Namely, vPRC1 and PRC2.2 are recruited to H2AK119ub1-modified nucleosomes through Rybp and Jarid2 respectively ^50-52^. Likewise, cPRC1 and PRC2 are targeted to nucleosomes containing H3K27me3 through Cbx or Eed respectively ^40, 41, 53^. In this study we found that enrichment of Rybp and H2AK119ub1 at PRC1 target promoter regions is increased during S and G2, indicating that polycomb repressive domains are dynamically assembled during interphase progression. This idea is further reinforced by the behaviour of PRC2 and H3K27me3, which previous reports have shown that also accumulates at target promoters during S and G2 phases ^19^, maybe through CDK2-mediated phosphorylation of Ezh2 ^26, 27^. Because the chromobox domain of Cbx7 can bind H3K27me3 ^54, 55^, accumulation of PRC2.2 at target genes during S and G2 phase ^19^ sustain our findings that recruitment of Cbx7-cPRC1 and DNA interactions among target promoters are enhanced during G2 compared to G1 phase. All PRC2 ^19^ and PRC1 repressive components analysed to date accumulate in G2 compared to G1, and thus, it might seem surprisingly that the catalytic PRC1 subunit Ring1b remain constant in G1 and G2 phases. However, Ring1b can bind to chromatin when partnered with Pcgf proteins ^56^ and Rybp is only required to enhance its ubiquitin transferase activity ^36, 37, 57^. Therefore, our results suggest that Ring1b interacts with Pcgf proteins in G1 phase to bind and moderately ubiquitinate H2AK119 at a large set of target genes. As interphase progresses, gradual recruitment of functionally efficient Rybp-containing vPRC1 complexes leads to enhanced ubiquitination of H2AK119 and formation of more effective repressive domains during S and G2 phases. We propose that initially low levels of H2AK119ub1 in G1 phase are expanded during S and G2 phases through positive PRC-dependent feedback loops that promote the formation of repressive chromatin domains highly enriched for PRC1 and PRC2 and associated histone modifications, in which stronger polycomb-mediated DNA interactions are established. Future studies will need to identify which are the molecular events that drive the accumulation of PRCs at target gene promoters upon G1 phase exit. Strikingly, Ezh2 ^26-29^ and other PRC1 and PRC2 subunits are phosphorylated during cell cycle transition (our analyses of datasets published in ^58^, data not shown). Thus, it is likely that phosphorylation of vPRC subunits by CDKs is a critical step required to coordinate polycomb repression with cell cycle progression.

Proliferating pluripotent cells need to maintain a delicate balance between preserving transcriptional memory during self-renewal and coordinating the erasure of such memory during induction of cell differentiation. This partly relies on the activity of chromatin factors, which are known to play decisive roles in the regulation of cell identity ^59^. Importantly, the standardized use of proliferating asynchronous populations of cells has perhaps led to the implicit assumption that the activity of epigenetic regulators is unaltered at different phases of the cell cycle. In this study, we have established that clearly this is not the case for the polycomb epigenetic system. We have provided solid evidence that PRC1 repression is diminished during the G1 phase to facilitate transcriptional induction of differentiation genes. However, this is probably only one side of the coin, because current evidence supports that PRCs endorse cells with the molecular memory required to perpetuate gene expression programs ^20, 21^, and therefore, we would expect that the molecular components of the polycomb system that encode transcriptional memory remain bound to target genes from G1 phase to mitosis. To identify such factors, future studies will need to analyse the distribution and function of PRCs on mitotic chromosomes. Overall, we propose that in proliferating stem cells, PcG proteins establish a self-perpetuating epigenetic cycle in which some components remain constant and provide the molecular memory, while others fluctuate to facilitate induction of cell differentiation in G1 phase.

Despite comprehensive evidence demonstrates that stem cells respond to developmental cues differently depending on the phase of the cell cycle in which they are found, very little is known about the molecular mechanisms involved ^5^. This is probably partly a consequence of existing technical limitations to carry out cell-cycle-phase-specific molecular and functional analysis. The FUCCI system ^60^ is an excellent tool to facilitate this task, and it has been used in pioneering studies that suggest that G1 phase-specific pluripotent cell differentiation is regulated by chromatin modifying enzymes ^16-19^. However, the underlying mechanisms remained largely unknown. Our findings reveal the molecular details as to how an epigenetic regulator can influence the ability of stem cells to induce differentiation depending on the phase of the cell cycle in which they are found. Our discoveries throw some light into the essential question as to why mESCs induce lineage transition preferentially during the G1 phase. Mitotic cell division is a major challenge for the maintenance of the epigenetic and transcriptional state in proliferating stem cells because it involves a breakdown of the nuclear envelope, chromosome condensation, drastic changes in PTMs and general downregulation of gene transcription ^61^. Therefore, in the subsequent G1 phase cells need to re-establish their cell-type-specific chromatin organization. We propose that differentiation signals are more effective at inducing transcriptional activation of differentiation genes in G1 cells, because at this phase of the cell cycle, they are reconstructing their chromatin organization, and they have temporarily lost a part of the mechanisms that reinforce maintenance of their cell identity (i.e. polycomb repressive domains highly enriched in H3K27me3 and H2AK119ub1).

Finally, it is worth mentioning that our results open new potential applications for currently developing inhibitors against CDKs ^62^, Ezh1/2 ^63^ and Ring1a/b ^64^, because they might be used individually or in combination to obtain populations of pluripotent cells enriched in G1 phase and/or with reduced polycomb repressive activity, and improve the efficiency of cell differentiation or nuclear reprogramming protocols.

## Methods

### Derivation of FUCCI-expressing mESC

Wild-type FUCCI-mESCs was described previously ^19^. FUCCI-expressing Ring1b inducible knockout (KO) mESCs were generated by introducing the FUCCI plasmid into previously derived *Ring1b* conditional KO mESCs (*Ring1a*^−/–^;*Ring1b*^fl/fl^;*Rosa26*::*CreERT2)* ^45^ (FUCCI-*Ring1b*^fl/fl^) using lipofectamine as previously described ^19^. FUCCI-expressing AID::Ring1b (AID::Ring1b;FUCCI) were obtained by transfecting the FUCCI plasmid into previously described Ring1a-/-; AID::Ring1b mESCs ^46^.

### Cell culture and flow cytometry sorting of FUCCI mESCs

Cells were grown on 0.1% gelatin-coated dishes in Dulbecco’s modified Eagle’s medium knockout (DMEM KO, Gibco) supplemented with 10% FBS, leukemia-inhibiting factor (LIF), penicillin/streptomycin (Biowest), L-glutamine (Biowest), 2-mercaptoethanol (Gibco) and hygromycin B (InvivoGen) at 37ºC and 5% CO_2_.

Deletion of Ring1b from FUCCI-Ring1b^fl/fl^ cell line was induced by treating mESCs with 800 nM 4-hydroxytamoxifen (Tmx, Sigma) for 48 hours and collected for downstream analyses 24 hours later (72 hours total). Auxin-induced degradation of Ring1b in *AID::Ring1b;FUCCI* was carried by adding water-dissolved auxin (IAA, Indole-3-acetic acid sodium salt, Sigma) to cell media to a final concentration of 500 µM as described previously ^46^. Cells were cell cycle sorted in an Aria Fusion flow cytometer as described previously ^19^ to obtain 1.5-2 million cells per cell cycle fraction for downstream analysis. Purity and cell cycle profile of sorted cell populations were routinely checked by propidium iodide (PI) staining (Table S1). To analyse the percentage of mitotic cells in each cell cycle fraction, sorted cells were DAPI stained and the number of cells in interphase or mitosis were determined using a fluorescence wide-field microscope. To check the percentage of cells that remains at each phase of the cell cycle, DNA content of cells was analysed by PI incorporation six hours after plating flow cytometry sorting in standard mESCs media.

### Retinoic acid differentiation of cell cycle-sorted FUCCI mESCs

For retinoic acid (RA)-induced differentiation analysis of gene expression, AID::Ring1b;FUCCI cells were cell cycle-sorted and plated at 2×10^5^ cells per well onto gelatin-coated six well tissue culture dish. Cells were cultured in complete ESC media without LIF and RA to a final concentration of 5µM. To induce the degradation of Ring1b, AID::Ring1b;FUCCI cells were treated with IAA at 500 µM final concentration. After six hours, cells were collected and subjected to mRNA analysis. For analysis of colony formation, AID::Ring1b;FUCCI cells were sorted and plated at low density (2000 cells per well onto gelatin-coated 6 well tissue culture dish) in complete DMEM KO medium containing 0.1 µM RA without LIF. To induce the degradation of Ring1b cells were treated with 500 µM IAA. Cells were exposed to this media for six hours and subsequently cultured in standard mESCs media in the presence of LIF and without RA nor IAA for five days. Cells were washed and fixed with paraformaldehyde (PFA) at final concentration of 4% in PBS for thirty seconds. Colonies with unequivocal differentiated morphology were quantified in four independent biological replicates.

### Chromatin immunoprecipitation followed by qPCR (ChIP-qPCR) or sequencing (ChIP-seq)

ChIP assays for Ring1b, Rybp, Cbx7 and H2AK119ub1 were performed as described previously (see table S2 and S3 for gene lists and reads coverage at gene promoter regions) ^19^. Briefly, 1.5-2 million cell-cycle sorted cells were fixed with 1% formaldehyde at room temperature in a rotating platform for 12 min. 2.5 µg antibodies per million cells were incubated with sonicated chromatin and in a rotating wheel at 4ºC overnight, followed by a five-hour incubation with Protein G magnetic beads (Dynabeads, Invitrogen). After reverse cross-link, immunoprecipitated DNA was ethanol-precipitated and resuspended in 200 µl of DNAse/RNAse free water. Libraries of immunoprecipitated DNA were generated from 1-5 ng of starting DNA with the NEBNext® Ultra DNA Library Prep for Illumina kit according to the manufacturer’s protocol at Centre for Genomic Regulation (CRG) Genomics Core Facility (Barcelona) and sequenced using HiSeq 2500 Illumina technology (Ring1b-MBL, Rybp, Cbx7 and H2AK119ub1 libraries) or NextSeq500 Illumina technology (Ring1b libraries). 20-30 million reads [50–base pair (bp) single reads] were obtained for each library.

Reads were aligned to the GENCODE NCBI m37 (mm9) genome using STAR 2.5.2 ^65^. Alignments with a quality score <200 were discarded by applying SAMtools 1.3.1 ^66^. BamCompare script from deepTools suite ^67^ was used to create bigwig files with the signal normalized by reads per million (RPM) and substracting the singal of the corresponding input sample. Peak calling was performed with MACS2 with following thresholds: Ring1b and Rybp (FDR<0.01), Ring1b-MBL, Ezh2 and Cbx7 (FDR<0.05), H2AK119ub1 (p-value<0.001) (Table S4). To calculate coverage (RPM) at promoters (−0.5 to +1.5 kb relative to TSS) CoverageView (Coverage visualization package for R. R package version 1.20.0.) was used. GraphPad software was used to represent normalized reads (RPM) for each analysed promoter as box plots and dot plots. Average binding plots around the TSS or peak centre were generated by counting normalized reads every 10 bp. Heatmap analyses of reads density in ChIP-seq experiments were performed by trimming RPM values between the minimum 5th percentile and the maximum 95th percentile. To compare different samples, genes were ranked according to G2. Gene Ontology analysis was performed using the Gene Ontology knowledge database (www.geneontology.org) and Enrichr (www.maayanlab.cloud/Enrichr).See table S2 for complete gene lists used. HC bivalent and hypermethylated promoters were previously defined ^19^.

ChIP-qPCR was performed using GoTaq qPCR Master Mix (Promega) with a StepOnePlus™ Real Time PCR system (Applied Biosystems). Enrichment was calculated relative to 1% input for each cycle cycle fraction. Details of antibodies and primers used are available in Table S4.

### mRNA sequencing and nascent RNA sequencing by 4sU-tagging

RNA was extracted with the RNeasy Plus Micro kit (Qiagen) according to manufacturer’s instructions and RNA concentration was measured using Qubit BR (Invitrogen). Libraries of mRNA (stranded) were generated from 300 ng of starting total RNA with the TruSeq stranded mRNA Library Prep kit for Illumina according to the manufacturer’s instructions at the Centre for Genomic Regulation (CRG) Genomics Core Facility (Barcelona) and sequenced using Nextseq 2000 Illumina technology. 25-30 million reads [50–base pair (bp) paired reads] were obtained for each library.

4sU-seq analyses (see table S3 for reads coverage at gene promoter regions) were carried out for cell cycle-sorted FUCCI-*Ring1b*^fl/fl^ cells as described previously ^19^. Briefly Tmx-treated and control cells were incubated during one hour at 37ºC with 4-thiouridine (Carbosynth) before flow cytometry sorting to obtain 1.5 million cells per fraction. Strand-specific RNA libraries were generated using 20 ng of 4sU-RNA and the TruSeq stranded mRNA Library Prep kit according to the manufacturer’s protocol without the initial polyA selection at Centre for Genomic Regulation (CRG) Genomics Core Facility (Barcelona) and sequenced using HiSeq2500 Illumina technology. 30-35 million reads [2×50+8– base pair reads] were obtained for each library. Analysis of previously generated 4sU-seq datasets ^19^ as well as mRNA-seq and 4sU-seq datasets generated in this study was performed as previously described ^19^. Average expression plots around TSS were generated by counting normalized reads (RPM) every 10 bp. Unsupervised clustering was carried out using TMM values in Cluster 3.0 and Java TreeViewer software. See table S2 for gene lists used.

### Capture-C

Capture-C analyses from G1 and G2 Tmx-treated and control cell cycle-sorted FUCCI-*Ring1b*^fl/fl^ cells were carried out as described previously with slight modifications ^68^. 2×10^6^ cells from each fraction were resuspended in 1.86 ml of complete medium. Cells were fixed with formaldehyde (1.89%) and incubated for 10 minutes on a rotating wheel at room temperature. The fixation reaction was stopped by adding cold glycine (final concentration 150 mM). Fixated cells were centrifuged (5 min 1000 rpm 4°C) and washed 1x with cold PBS. After centrifugation, cells were resuspended in cold lysis buffer (10 mM Tris pH 8, 10 mM NaCl, 0.2% NP-40, 1x protease inhibitors (Roche)) and incubated for 20 min on ice. Subsequently, the pellet was centrifuged (5 min, 1800 rpm, 4°C) and resuspended in 200 μl of lysis buffer. Samples were frozen for subsequent experimental procedures at -80ºC. Lysates were thawed on ice, pelleted, and resuspended in 1x DpnII buffer (New England Biology). The lysates were incubated with 0.28% final concentration of SDS (Thermo Fischer Scientific) in 200 µl reaction volume for 1hr at 37°C in a thermomixer shaking at 500rpm (30 sec on/off). Reactions were quenched with 1.67% final concentration of Triton X-100 for 1hr at 37°C in a thermomixer shaking at 500rpm (30 sec on/off) and digested for 24 hours with 3×10µl DpnII (produced in-house) at 37°C in a thermomixer shaking at 500rpm (30 sec on/off). Digested chromatin was ligated with 8 µl T4 Ligase (240U, Themo Firsher Scientific) for 20 hours at 16°C. The nuclei containing ligated chromatin were pelleted to remove any non-nuclear chromatin, reverse cross-linked and the ligated DNA was phenol-chloroform purified. Samples were resuspended in 100 µl water and sonicated for 135 sec using Covaris ME220 with microTUBE-50 AFA tubes to achieve a fragment size of approximately 200bp. Two reactions of 2-4 µg DNA each were adaptor-ligated and indexed using NEBNext Ultra II DNA Library Prep Kit for Illumina (New England Biolabs) and NEBNext Multiplex Oligos for Illumina (New England Biolabs). The libraries were amplified with 7 PCR cycles using Herculase II Fusion Polymerase kit (Agilent). Libraries hybridisation was performed as described previously ^68^ using probes mapping to 122 polycomb target genes promoters and 78 control active gene promoters. Libraries were performed in biological quadruplicates and sequenced using illumina NextSeq 500.

Paired-end reads were aligned to mm10 and filtered for Hi-C artefacts using HiCUP ^69^ and Bowtie 2 ^70^, with fragment filter set to 100-800bp. Counts of reads aligning to captured gene promoters and interaction scores (significant interactions) were then called by CHiCAGO ^71^. For visualisation of Capture-C data, weighted pooled read counts from CHiCAGO data files were normalized to total read count aligning to captured gene promoters in the sample and further to the number of promoters in the respective capture experiment and multiplied by a constant number to simplify genome-browser visualization, using the following formula: normCounts=1/cov*nprom*100000. Bigwig files were generated from these normalized read counts. For scatter and bar plots analyses we first determined interactions between promoters and regions that were accessible by ATAC analysis (ATAC peaks) using CHiCAGO (cutoff score >= 5). Next, for each promoter-ATAC peak interaction, we quantified the sum of normalized read counts or CHiCAGO scores across all DpnII fragments overlapping this interval. For metaprofiles, these counts were further normalized to the counts within the DpnII fragment overlapping the interaction peak summit in the G2 sample. Reads were moreover flipped to represent directional profiles from promoter-distal (−40) to promoter-proximal (+40) reads. Ring1b and Cbx7-bound promoters were filtered using previously published ChIP-seq data ^34^.

### 4C-seq

Analysis of chromatin interactions between *Six2* and *Six3* promoters was carried out as described previously in ^72^ from G1 and G2-sorted FUCCI-Ring1b^fl/fl^ cells. For each condition 4 million cells were used as starting material. The resulting DNA products were amplified by inverse PCR using the Expand Long Template PCR system (Roche) with primers designed within the *Six2* promoter. Primers used are available in table S4. 4C samples were sequenced on an Illumina HiSeq 2500 sequencer, generating individual reads of 74 bp at Center for Biomics Erasmus University Medical Center in Rotterdam (The Netherlands). 4C-seq data analysis was performed as described in ^72^.

### Western blot and cell fractionation

Western blot of whole cell extracts and cell fractionation were carried out as previously in ^19^. One million sorted FUCCI-mESCs were used for each condition. Equivalent amount of protein was loaded for whole cell extracts western blot, while the same number of cells were loaded in cell fractionation experiments. Quantification of band intensity and normalization against LaminB and H3 was carried out using ImageJ.

### Statistical analysis

R 3.5.1 was used to carry out statistical analyses. In boxplots, whiskers denote the interval within 1.5× the interquartile range, and P values were calculated using Mann-Whitney test (significant differences: *P<0.05, **P<0.01, ***P<0.001, ****P<0.0001). Statistical analysis of average mapped reads around the TSS between G1, S and G2 samples was carried out using analysis of variance (ANOVA), comparing all samples in a window of −0.5 to +1.5 kb from TSS (significant differences P < 0.0001).

## Data access

Datasets are available at GEO-NCBI with accession number GSE207997 with a private token that will be released upon publication acceptance.

## Author contributions

DL designed and conceptualized the study. DL and HGA designed experiments. HGA, ME, LLO, MAF, AG, ED and TP performed and analysed experiments. JMM and AF performed bioinformatic analyses. PCS, ASP, ARI and RJK provided scientific advice and resources. DL obtained funding and supervised research.

## Acknowledgments

We thank core facilities at GENYO for excellent technical support. We also thank the genomics unit at the CRG for assistance with RNA-seq and ChIP-seq experiments.

## Figure legends

**Figure S1.**
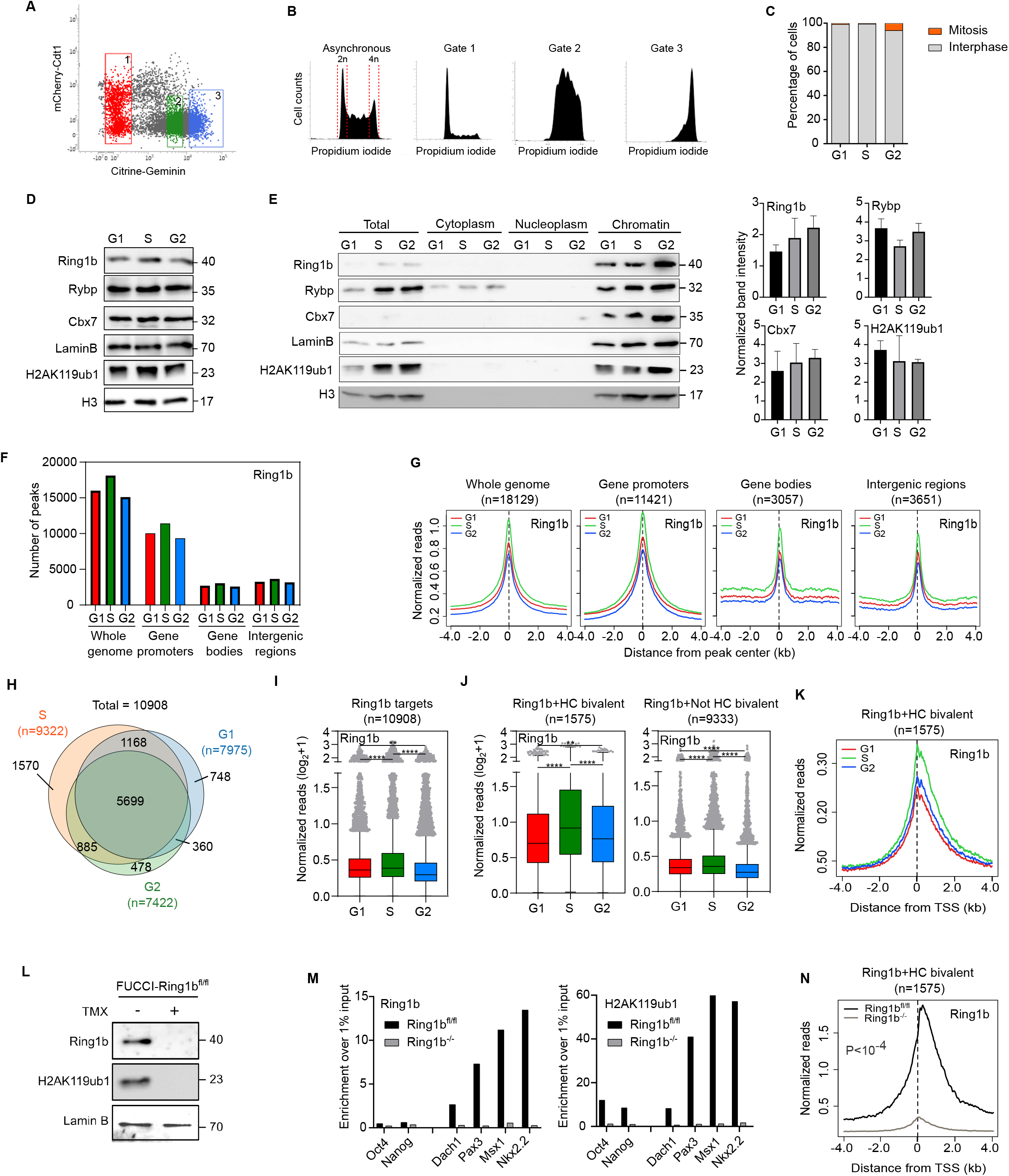
The global amount of PRC1 subunits on chromatin is similar during G1, S and G2 phases. **(A)**Flow cytometry dot plot analysis of wild-type FUCCI-mESCs (expressing mCherry-Cdt1 and Citrine-Geminin fusion proteins) indicating the sorting gates used to obtain cell populations enriched in G1 (Gate 1), S (Gate 2) and G2 (Gate 3) phases of the cell cycle. **(B)**Flow cytometry analysis of DNA content in cells stained with propidium iodide after sorting as indicated in (A). **(C)**Percentage of cells in interphase (grey) or mitosis (orange) at indicated fractions after cell cycle sorting as assayed by fixation, DAPI staining and microscopy analysis. **(D)**Whole cell lysate western blots of wild-type FUCCI-mESCs comparing the level of Ring1b, Rybp, Cbx7 and H2AK119ub1 in G1, S and G2 phases. Lamin B and histone 3 (H3) were used as a loading control. Molecular weight is indicated in kDa. **(E)**Western blot analyses of Ring1b, Rybp, Cbx7 and H2AK119ub1 proteins upon biochemical cell fractionation of wild-type FUCCI-mESCs in the indicated cell-cycle phases. Lamin B and histone 3 (H3) were used as loading control for the chromatin fraction. Molecular weight is indicated in kDa. Histograms showing quantification of the intensity of the bands in the chromatin fraction (using Lamin B and H3 as normalizers) is shown on the right panel. Mean ± SEM of three experiments is shown. **(F)**Histogram showing the number of peaks detected in the Ring1b ChIP-seq across the whole genome, gene promoters (±2 kb from TSS), gene bodies and intergenic regions at indicated cell cycle phases. **(G)**Average signal of Ring1b binding at peaks detected in S phase and categorized as in (F) in G1 (red), S (green) and G2 (blue) phases. **(H)**Venn diagram showing the overlap between Ring1b target promoters detected in G1, S and G2 phases. Total number of promoters bound by Ring1b at any phase of the cell cycle is indicated (n=10908). **(I, J)** Quantification of Ring1b-binding signal at the promoter region (−0.5 kb to +1.5 kb relative to TSS) of target genes (I) that are HC bivalent or not (J) (as identified in figure 1D) at the indicated cell cycle phases. Asterisks (*) mark statistically significant differences. **(K)** Average binding profile of Ring1b around the TSS of HC bivalent Ring1b target promoters in G1 (red), S (green) and G2 (blue) phases using an alternative anti-Ring1b antibody (MBL D139-3) for ChIP-seq. **(L)**Western blots of whole cell extracts of Ring1b and H2AK119ub1 in FUCCI-Ring1b^fl/fl^ mESC before (−) and after (+) 72h Tmx treatment. Lamin B was used as loading control. Molecular weight is indicated in kDa. **(M)**Analysis by ChIP-qPCR comparing the enrichment of Ring1b (left) and H2AK119ub1 (right) at candidate PRC1-target promoter regions (Dach1, Pax3, Msx1, Nkx2-2) in an asynchronous population of parental FUCCI-Ring1b^fl/fl^ and 72h Tmx-treated Ring1b^-/-^. Active (Oct4, Nanog) gene promoters were used as negative controls. **(N)**Average binding signal of Ring1b around the TSS of HC bivalent target promoters in parental FUCCI-Ring1b^fl/fl^ (black line) and 72h Tmx-treated Ring1b^-/-^ (grey line) mESCs. *P* indicates Mann-Whitney test p-value.

**Figure S2.**
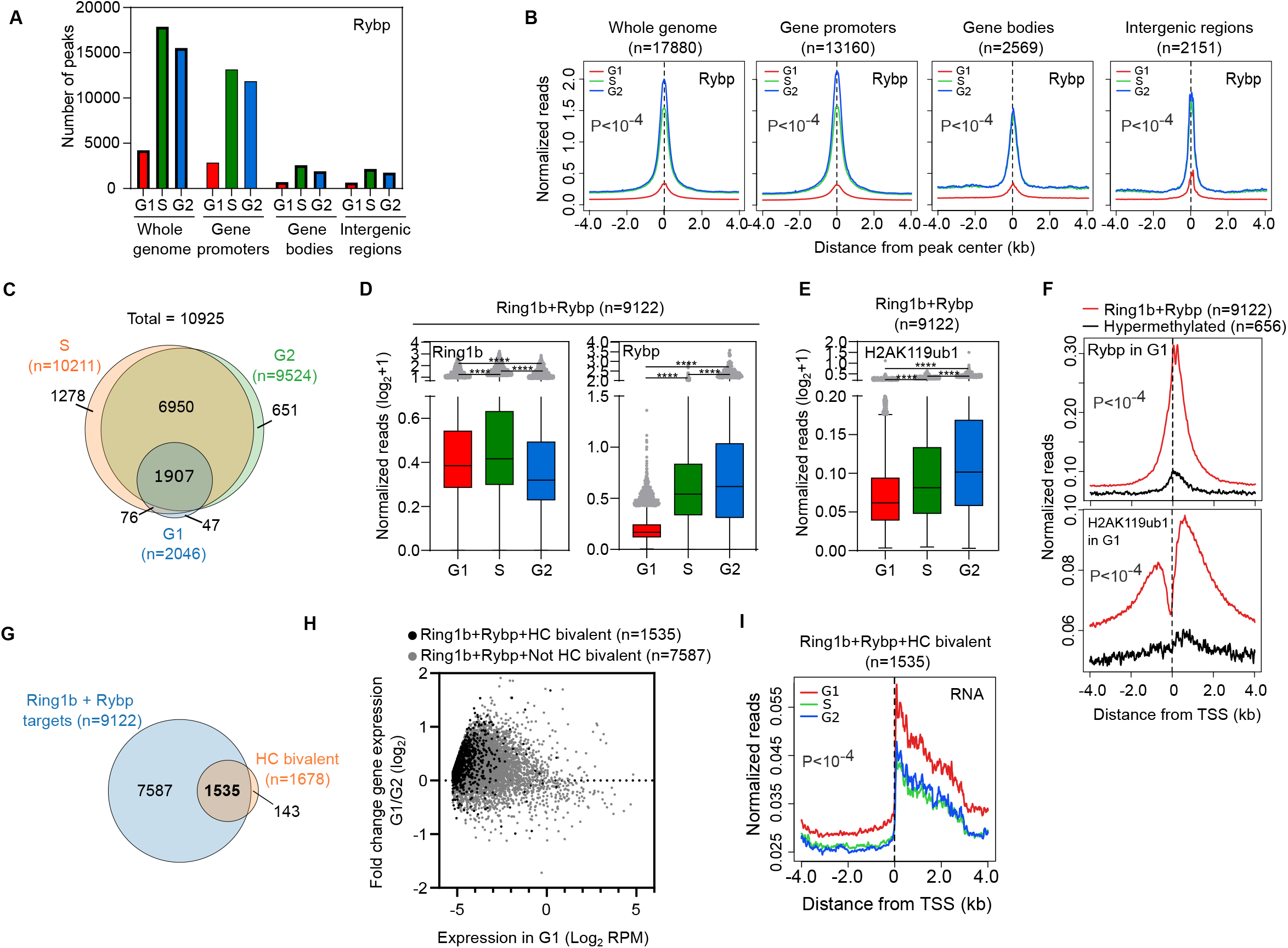
Rybp and H2AK119ub1 are recruited to target promoters during G1 phase in mESCs. **(A)**Histogram showing the number of peaks detected in the Rybp ChIP-seq across the whole genome, gene promoters (±2 kb from TSS), gene bodies and intergenic regions at indicated cell cycle phases. **(B)**Average signal of Rybp binding at peaks detected in S phase and categorized as in (F) in G1 (red), S (green) and G2 (blue) phases. *P* indicates ANOVA test p-value. **(C)**Venn diagram showing the overlap between Rybp target promoters detected in G1, S and G2 phases. Total number of promoters bound by Rybp at any phase of the cell cycle is indicated (n=10925). **(D, E)** Quantification of Ring1b, Rybp (D) and H2AK119ub1 (E) binding signal at the promoter region (−0.5 kb to +1.5 kb relative to TSS) of Ring1b and Rybp targets in the indicated phases of the cell cycle. Asterisks (*) mark statistically significant differences. **(F)**Average enrichment profile of Rybp (top) and H2AK119ub1 (bottom) around the TSS at Ring1b and Rybp target promoters (red) compared to hypermethylated promoters (black) in G1 phase. Hypermethylated promoters were previously defined ^19^. *P* indicates Mann-Whitney test p-value. **(G)**Venn diagram showing the overlap between HC bivalent promoters (orange) and Ring1b+Rybp target promoters (red). HC bivalent promoters were previously defined ^19^. **(H)**MA plot showing fold change gene expression between cells in G1 and G2 phases (4sU-seq normalized reads mapping from the TSS to +3kb) at HC bivalent Ring1b+Rybp targets (black dots) and Not HC bivalent Ring1b+Rybp targets (grey dots). Nascent RNA expression in G1 is represented in the x-axis. **(I)**Average 4sU-seq RNA reads at Ring1b+Rybp+HC bivalent promoters in G1 (red), S (green) and G2 (blue) phases. *P* indicates ANOVA test p-value.

**Figure S3.**
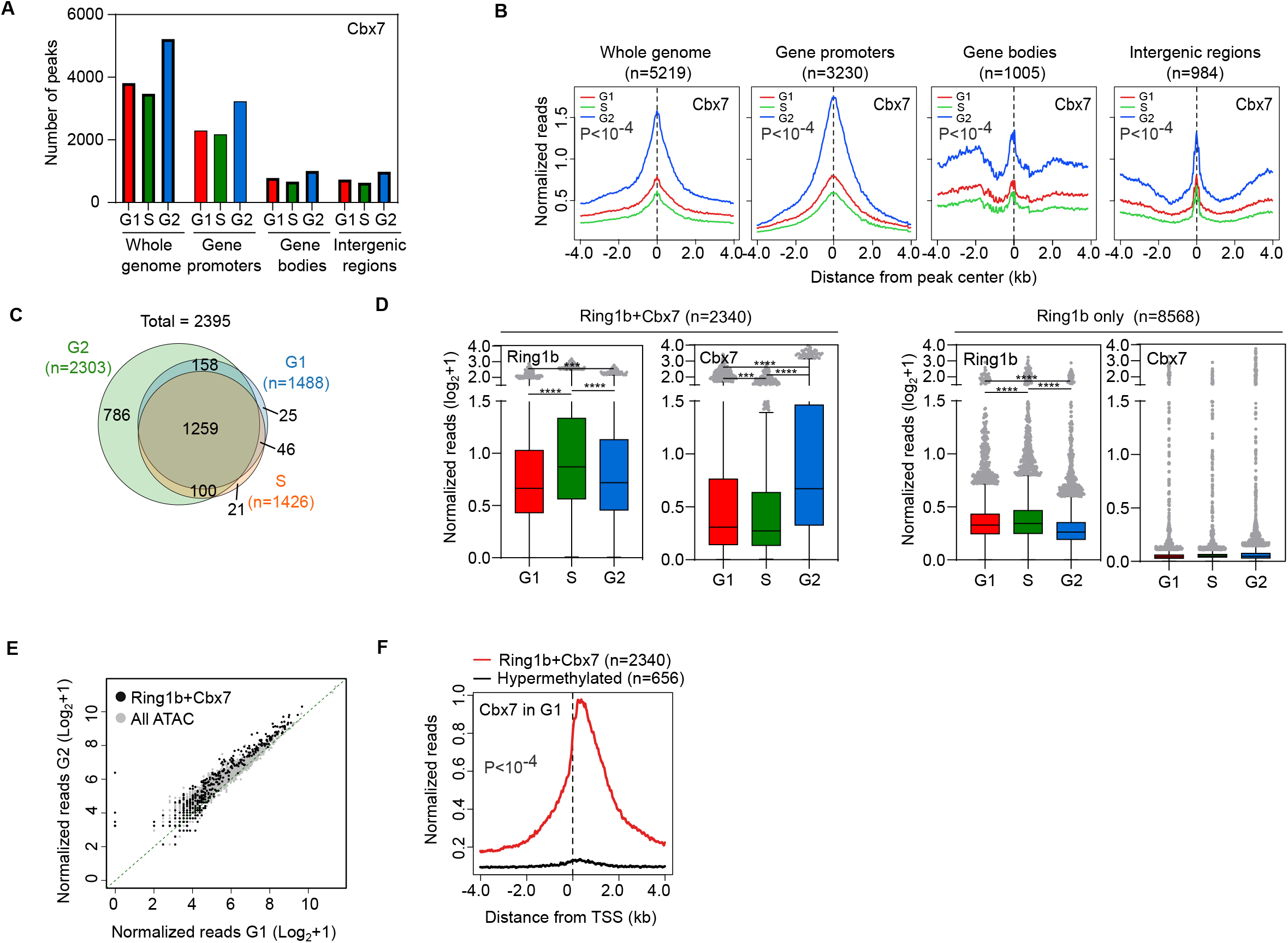
Cbx7 is recruited to target promoters during G1 phase in mESCs. **(A)**Histogram showing the number of peaks detected in the Cbx7 ChIP-seq across the whole genome, gene promoters (±2 kb from TSS), gene bodies and intergenic regions at indicated cell cycle phases. **(B)**Average signal of Cbx7 binding at peaks detected in S phase and categorized as in (F) in G1 (red), S (green) and G2 (blue) phases. *P* indicates ANOVA test p-value. **(C)**Venn diagram showing the overlap between Cbx7 target promoters detected in G1, S and G2 phases. Total number of promoters bound by Cbx7 at any phase of the cell cycle is indicated (n=2395). **(D)**Quantification of Ring1b and Cbx7 binding signal at the promoter region (−0.5 kb to +1.5 kb relative to TSS) of Ring1b-bound promoters that are targeted by Cbx7 (left panel) or not (right panel) at the indicated phases of the cell cycle. Asterisks (*) mark statistically significant differences. **(E)**Scatter plot showing the Capture-C signal in G1 (x axis) compared to G2 (y axis) of interactions (n=2034) taking place between accessible promoters (n=179) and other regions of the genome (black and grey dots). Black dots highlight interactions (n=392) involving Ring1b+Cbx7-bound promoters (n=39). **(F)**Average enrichment profile of Cbx7 around the TSS at Ring1b+Cbx7 target promoters (red) compared to hypermethylated promoters (black) in G1 phase. Hypermethylated promoters were previously defined ^19^. *P* indicates Mann-Whitney test p-value.

**Figure S4.**
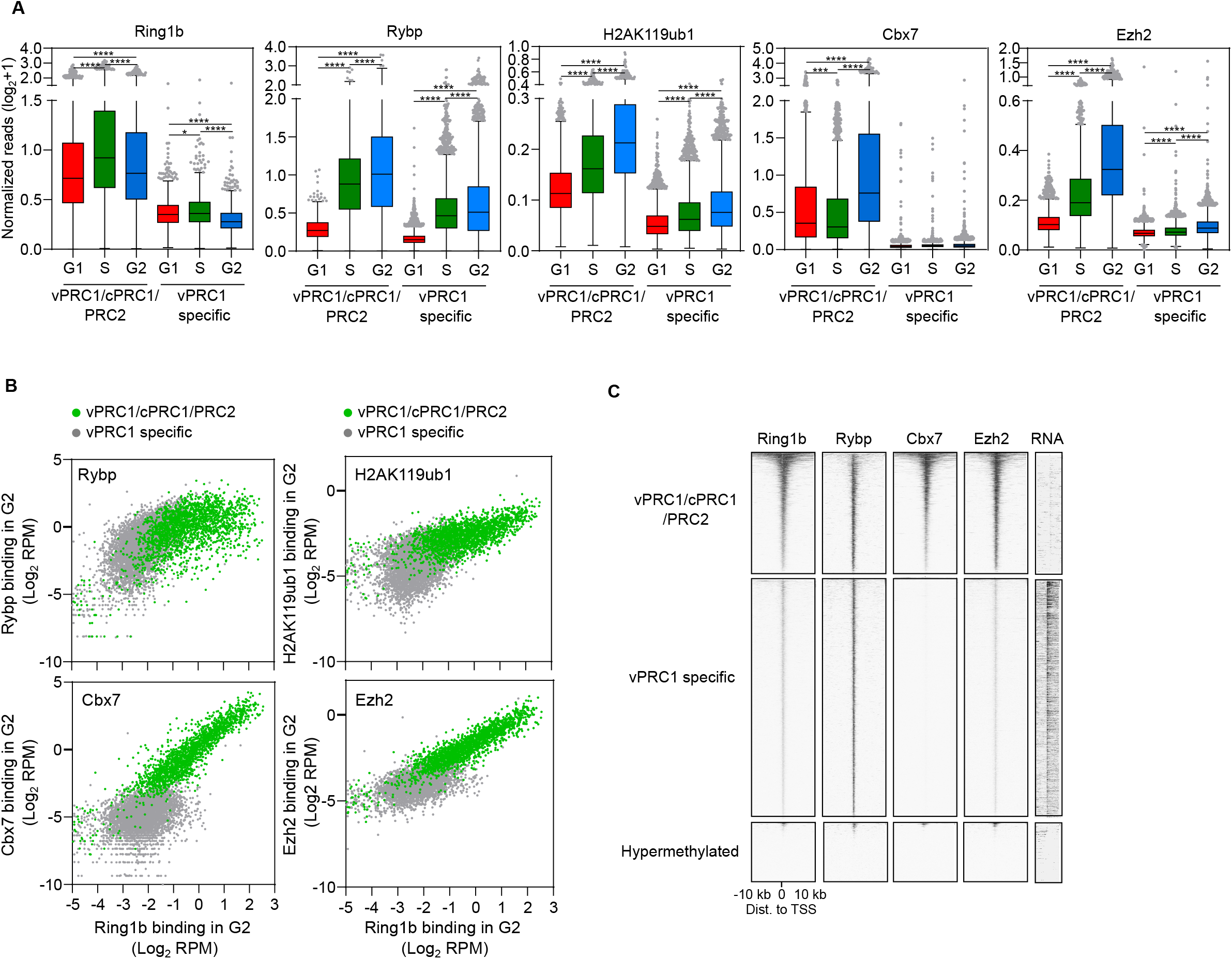
vPRC1/cPRC1/PRC2 target genes display higher levels of Ring1b, Rybp, Cbx7, Ezh2 and H2AK119ub1 binding than vPRC1 specific target genes. **(A)**Quantification of Ring1b, Rybp, H2AK119ub1, Cbx7 and Ezh2 binding signal at the promoter region (−0.5 kb to +1.5 kb relative to TSS) of vPRC1/cPRC1/PRC2 and vPRC1 specific target promoters in the indicated cell cycle phases. Asterisks (*) mark statistically significant differences. **(B)**Correlation analysis between the binding signals (−0.5 kb to +1.5 kb relative to TSS) of Rybp, H2AK119ub1, Cbx7 or Ezh2 compared to Ring1b in G2 phase at vPRC1/cPRC1/PRC2 (green dots) and vPRC1 specific (grey dots) target promoters. **(C)**Heatmaps of normalized Ring1b, Rybp, Cbx7 and Ezh2 ChIP-seq reads and RNA expression around the TSS (±10 kb) of vPRC1/cPRC1/PRC2, vPRC1 specific and control hypermethylated promoters in G2 phase. Genes are ranked according to the signal of Ring1b binding.

**Figure S5.**
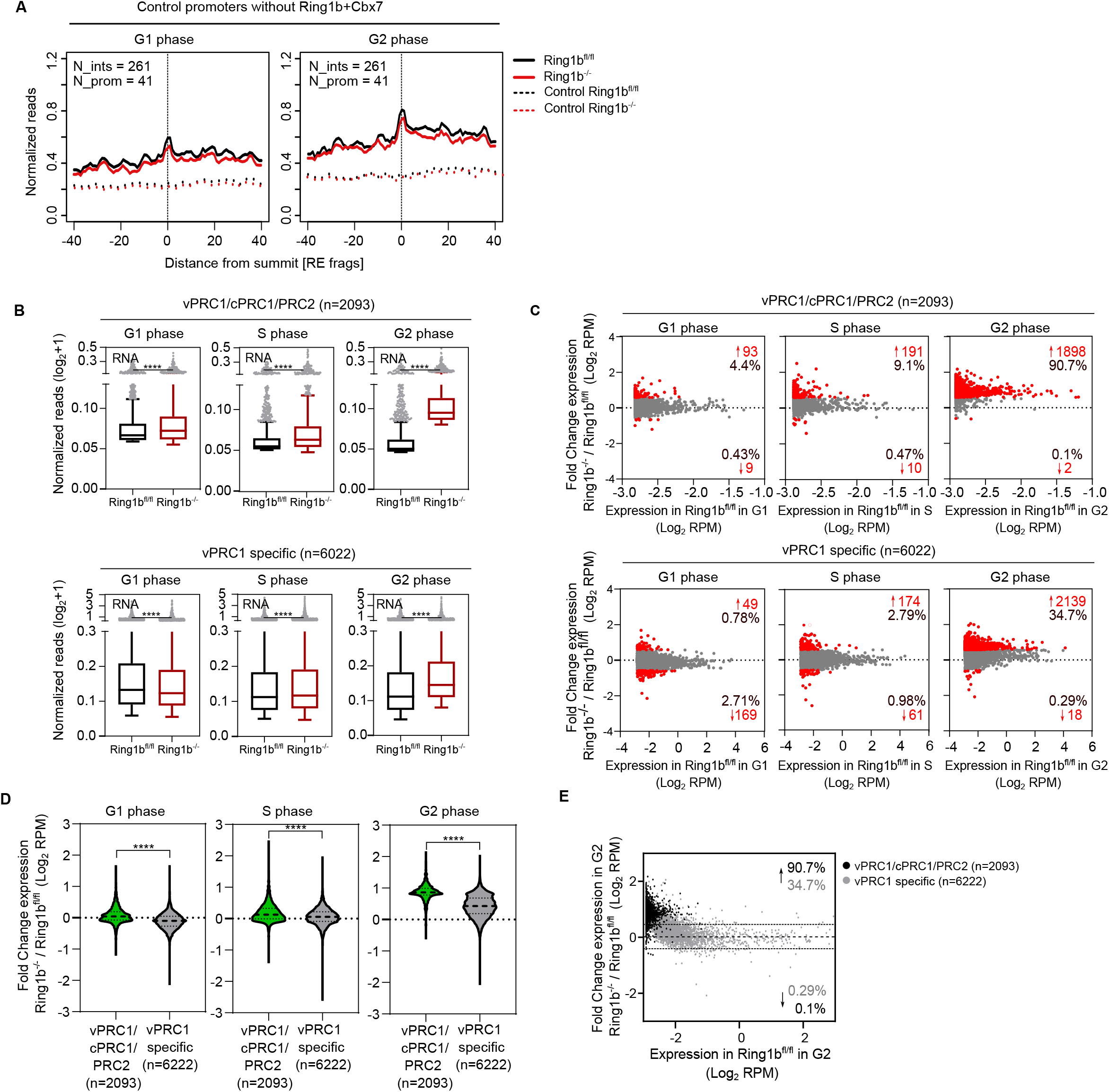
Depletion of Ring1b barely perturbs the transcriptional repression of target genes in G1 phase. **(A)**Average normalized reads of 3D chromatin interactions (number of interactions = 261) measured by Capture-C of control promoters that are not bound by Ring1b+Cbx7 involving at least one promoter (number of promoters=41) in Ring1b^fl/fl^ (black line) and Ring1b^-/-^ (red line) cells during G1 (left) and G2 (right) phases. X axis represents the distance from the summit of the interaction peak measured as a function of DpnII restriction fragments. Dashed lines show enrichment at distance-matched control sites from each promoter and interaction in the opposite direction. **(B)**Boxplot of nascent RNA reads mapped to the proximal promoter region (TSS to +3Kb) of vPRC1/cPRC1/PRC2 promoters (top panel) and vPRC1 specific targets (bottom panel) in Ring1b^fl/fl^ and Ring1b^-/-^ cells in G1 (left), S (middle) and G2 (right) phases. Asterisks (*) mark statistically significant differences. **(C)**MA plot showing fold changes in nascent RNA between Ring1b^-/-^ and Ring1b^fl/fl^ cells at vPRC1/cPRC1/PRC2 (top panel) and vPRC1 specific (bottom panel) target genes during G1, S and G2 phases. Genes showing a fold change > 1.5 are represented as red dots. Number and percentage (relative to total group targets) of deregulated genes are indicated. **(D)**Violin plots showing the fold change of nascent RNA mapped to the proximal promoter region (TSS to +3kb) in Ring1b^-/-^ and Ring1b^fl/fl^ cells at vPRC1/cPRC1/PRC2 targets (green) and vPRC1 specific (grey) target genes in G1, S and G2 phases. Asterisks (*) mark statistically significant differences. **(E)**MA plot showing fold changes of nascent RNA mapped to the proximal promoter region (TSS to +3kb) between Ring1b^-/-^ and Ring1b^fl/fl^ cells at vPRC1/cPRC1/PRC2 (black dots) and vPRC1 specific (grey dots) target genes relative to gene expression in Ring1b^fl/fl^ cells in G2 phase. Percentage of genes up or down-regulated (above threshold FC>1.5) are shown.

**Figure S6.**
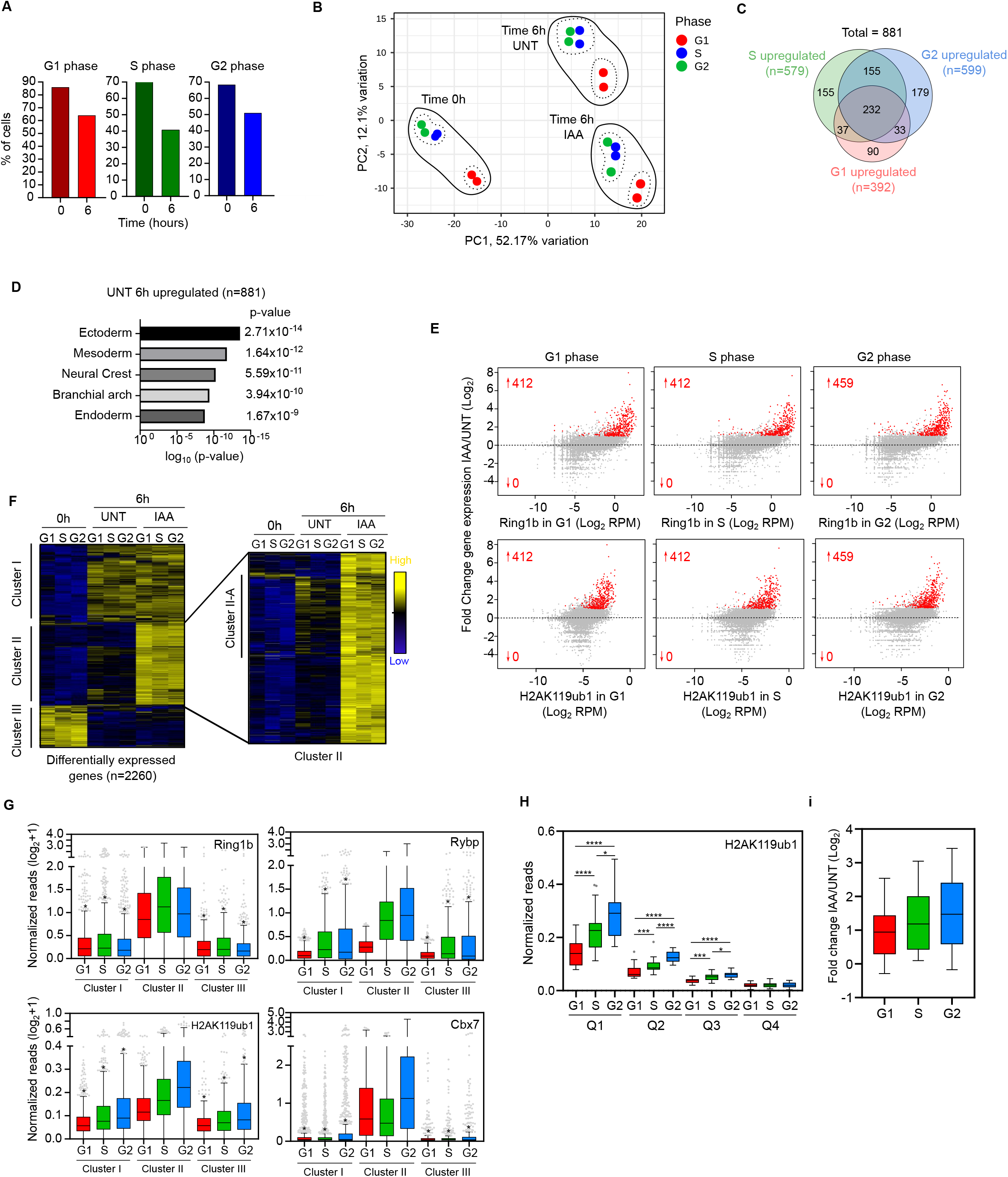
Treatment with retinoic acid induces the expression of ectoderm genes. **(A)**Histogram showing the percentage of AID::Ring1b;FUCCI cells in each phase of the cell cyle right (0h) or after six hours of (6h) flow cytometry sorting in G1, S or G2 phases. Cells were fixed and stained with propidium iodide to determine the phase of the cell cycle. **(B)**Principal component analysis of gene expression of AID::Ring1b;FUCCI cells that had been cell cycle sorted (time 0h) and treated with RA for 6h. Ring1b-depleted (time 6h, IAA) and control (time 6h UNT) cells are shown. Two biological replicates per experimental condition are represented. **(C)**Venn diagram showing the overlap of 881 genes upregulated (FC>2, FDR<0.05) after 6h or RA treatment in cells sorted in the three different phases of interphase. **(D)**Tissues gene ontology analysis of genes upregulated (FC>2, FDR<0.05, n=881) after 6h of RA treatment (identified in figure S6C). **(E)**MA plot showing the fold changes in gene expression (TMM) between IAA-treated (IAA) and untreated (UNT) cells 6h upon RA stimulation. X-axis shows the enrichment of Ring1b (top panel) or H2AK119ub1 (bottom panel) at the promoter (−0.5+1.5kb from TSS) of annotated genes in the mouse genome. Number of significantly up or down-regulated genes are indicated (FC>2, FDR<0.05). **(F)**Hierarchical clustering analysis of RNA expression levels (TMM) of the summation of differentially expressed genes (0h vs 6h UNT and 0h vs 6h IAA) at any cell cycle phase during RA induction. Cluster I highlight genes responding to RA stimulation but not to Ring1b depletion. Cluster II includes genes that are de-repressed upon Ring1b depletion. These include a group of genes that are transcriptionally induced by RA preferentially during the G1 phase (cluster II-A). Cluster III contains genes downregulated during RA differentiation. **(G)**Boxplot showing the enrichment of indicated PRC1 proteins at the promoter region (−0.5 kb to +1.5 kb relative to TSS) of genes belonging to the three clusters identified in figure S6F. Asterisks (*) mark statistically significant differences (p<0.0001) compared to the same cell cycle phase of cluster II. **(H)**Boxplot showing H2AK119ub1 enrichment at the promoter region (−0.5 kb to +1.5 kb relative to TSS). Genes preferentially upregulated in G1 compared to G2 (described in figure 6C) were classified into quartiles based on H2AK119ub1 enrichment at G2 phase. Asterisks (*) mark statistically significant differences. **(I)**Boxplots showing the fold change of RNA expression (TMM) between IAA-treated and untreated cells at 6h after RA treatment in cell cycle sorted cells.

## Table legends

Table S1.Purity check of cell cycle–sorted FUCCI-mESCs used in this study.

Table S2. List of genes used in this manuscript.

Table S3.ChIP-seq enrichment and expression values.

Table S4.Reagents and published datasets used in this manuscript.

